# Neuronal perception of the social environment intergenerationally controls germline development and generation time in *C. elegans*

**DOI:** 10.1101/2020.09.03.279158

**Authors:** Marcos Francisco Perez, Mehrnaz Shamalnasab, Alejandro Mata-Cabana, Simona Della Valle, María Olmedo, Mirko Francesconi, Ben Lehner

## Abstract

An old and controversial question in biology is whether information perceived by the nervous system of an animal can ‘cross the Weismann barrier’ to alter the phenotypes and fitness of their progeny. Here we show that such intergenerational transmission of sensory information occurs in the model organism, *C. elegans*, with a major effect on fitness. Specifically, that perception of social pheromones by chemosensory neurons controls the post-embryonic timing of development of one tissue – the germline – relative to others in an animal’s progeny. Neuronal perception of the social environment thus intergenerationally controls the generation time of this animal.

## INTRODUCTION

Phenotypic variation within a population – whether behavioral, morphological or physiological – is the basis of individuality and the raw material upon which natural selection acts. While phenotypic variation can arise due to genetic variation, the environment can play a large role: frequently a single genotype can produce alternative phenotypes in response to different environments – an ability called phenotypic plasticity (Fusco and Minelli 2010). The biotic environment, such as chemical or social cues from conspecifics or predators can also be an important source of variation. For example, *Caenorhabditis elegans* nematode worms growing in an unfavorable environment of high population density and temperature and low food enter dauer diapause, an alternative larval stage characterized by morphological, physiological and behavioral alterations and longevity (Golden and Riddle 1984). *C. elegans* and other nematodes sense population density by secreted pheromones called ascarosides, analogous to quorum sensing in bacteria (Choe et al. 2012). Although dauer entry is the most drastic transition induced by ascaroside pheromones, a range of plastic phenotypic responses are observed at various stages of the worm lifecycle, from slowed development (Ludewig et al. 2017) to shifts in reproductive traits, metabolism and social behavior (McGrath and Ruvinsky 2019).

Phenotypic plasticity can also sometimes operate across generations, whereby parental or ancestral environment induces phenotypic changes in progeny. For example, exposure of the water flea *Daphnia cucullata* to chemical cues indicating the presence of predators results in the production of progeny born with larger defensive helmets (Agrawal et al. 1999) and female population density reduces larval size in the coral reef fish *Pomacentrus amboinensis* (McCormick 2006). Intergenerational regulation of phenotypic plasticity has also been reported in mammals, with parental perception of higher population density in the red squirrel *Tamiasciuris hudsonicus* accelerating progeny growth rates (Dantzer et al. 2013).

While the control of somatic developmental timing has been well studied in *C. elegans* and *Drosophila melanogaster* (Thummel 2001), little is known about how the coordination of developmental timing between tissues changes in response to environmental cues. In *C. elegans* the timing of somatic and germline developmental can vary across individuals and genotypes (Kawasaki et al. 2007, Perez et al. 2017, Poullet et al. 2016) but the study of soma/germline developmental coordination has been mostly limited to the ongoing bidirectional communication that links germline development with that of the functionally and physically intertwined somatic gonad (Gilboa and Lehmann 2006, Korta and Hubbard 2010). In both *C. elegans* and *Drosophila*, neuronal signaling regulates germline proliferation in response to environmental conditions (Dalfó et al. 2012, LaFever and Drummond-Barbosa 2005). However, these reports of neuronal signaling to the germline are limited to a single generation.

*C. elegans* has emerged as a powerful model organism for studying intergenerational and transgenerational epigenetic inheritance (Perez and Lehner 2019, Baugh and Day 2020). Although numerous examples of intergenerational and transgenerational epigenetic inheritance have now been documented in *C. elegans*, most of these involve the stable or transient inheritance of RNAi- or piRNA-triggered gene silencing (Ashe et al. 2012, Bagijn et al. 2012, Luteijn et al. 2012, Rechavi et al. 2011, Shirayama et al. 2012, Beltran et al, 2020), the inheritance of environmentally-triggered changes in small RNA abundance (Rechavi et al. 2014, Schott et al. 2014) or of environmentally-triggered changes in repeat and transgene expression (Klosin et al. 2017). It remains unclear how most of these phenomena relate to physiological intergenerational plastic responses to natural environments (Perez and Lehner 2019). However, recent reports of multigenerational effects of the environment on stress resistance and life history traits (Burton et al. 2017, Hibshman et al. 2016, Jobson et al. 2015, Webster et al. 2018, Palominos et al., 2017), on behavioral avoidance of pathogenic bacteria (Kaletsky et al. 2019, Moore et al. 2019) and on pathogen resistance (Burton et al., 2020) suggest that non-genetic inheritance in *C. elegans* may indeed have ecological significance.

In general, though, the intergenerational responses to environmental perception in this and other organisms remain unclear. Indeed, whether the effects of stimuli perceived in somatic tissues – and in particular by the nervous system – are communicated to the germline across the ‘Weismann barrier’ - remains largely unknown, with only a few examples of the transmission of environmental information perceived by neurons to future generations (Dias and Ressler 2014, Gapp et al. 2014a, Gapp et al. 2014b, Remy 2010), even though the capacity to transmit information from neurons to the germline likely exists (Devanapally et al. 2015, Posner et al. 2019).

We previously reported that maternal age is an important cause of phenotypic variation in *C. elegans* progeny (Perez et al. 2017). Most of these phenotypes, including size at hatching, time to reach adulthood and starvation resistance, could be explained by an age-dependent increase in the supply of a lipoprotein complex, vitellogenin, to oocytes. However, we also reported that maternal age affects the relative developmental timings of somatic and germline tissues and that this effect was independent of embryonic vitellogenin levels. This indicated that somatic and germline developmental timings are partially uncoupled in isogenic individuals, and that their relative timing can be altered by changes in maternal physiology.

Here we report that the timing of post-embryonic germline development in isogenic animals in a constant environment is actually controlled by neuronal perception of the social environment by an animal’s parent. Neuronal perception in one generation – in this case, of a social cue – can therefore control the timing of development of a tissue within an animal in the next generation. The nervous system therefore communicates information about the social environment of an animal to its progeny to determine their generation time.

## RESULTS

### Exposure of parents to pheromone induces germline delay in progeny

We previously described in *C. elegans* an effect of maternal age on the relative timings of somatic and germline maturation in progeny (Perez et al. 2017). Although progeny of the youngest mothers reach both somatic and germline maturity later, this delay impacts somatic timing disproportionately, leading to the production of embryos sooner after the adult molt. During the course of these experiments, we noticed that if parental worms were transferred to a fresh plate during adulthood (Fig 1a), their progeny reached germline maturity more rapidly than worms whose parents that remained on the same, previously used plate (Fig 1b, c). While time to reach somatic maturity varied, time to germline maturity was robustly and consistently reduced with respect to the adult molt, and nearly always reduced in absolute terms, impacting the progeny minimum generation time.

**Figure 1.**
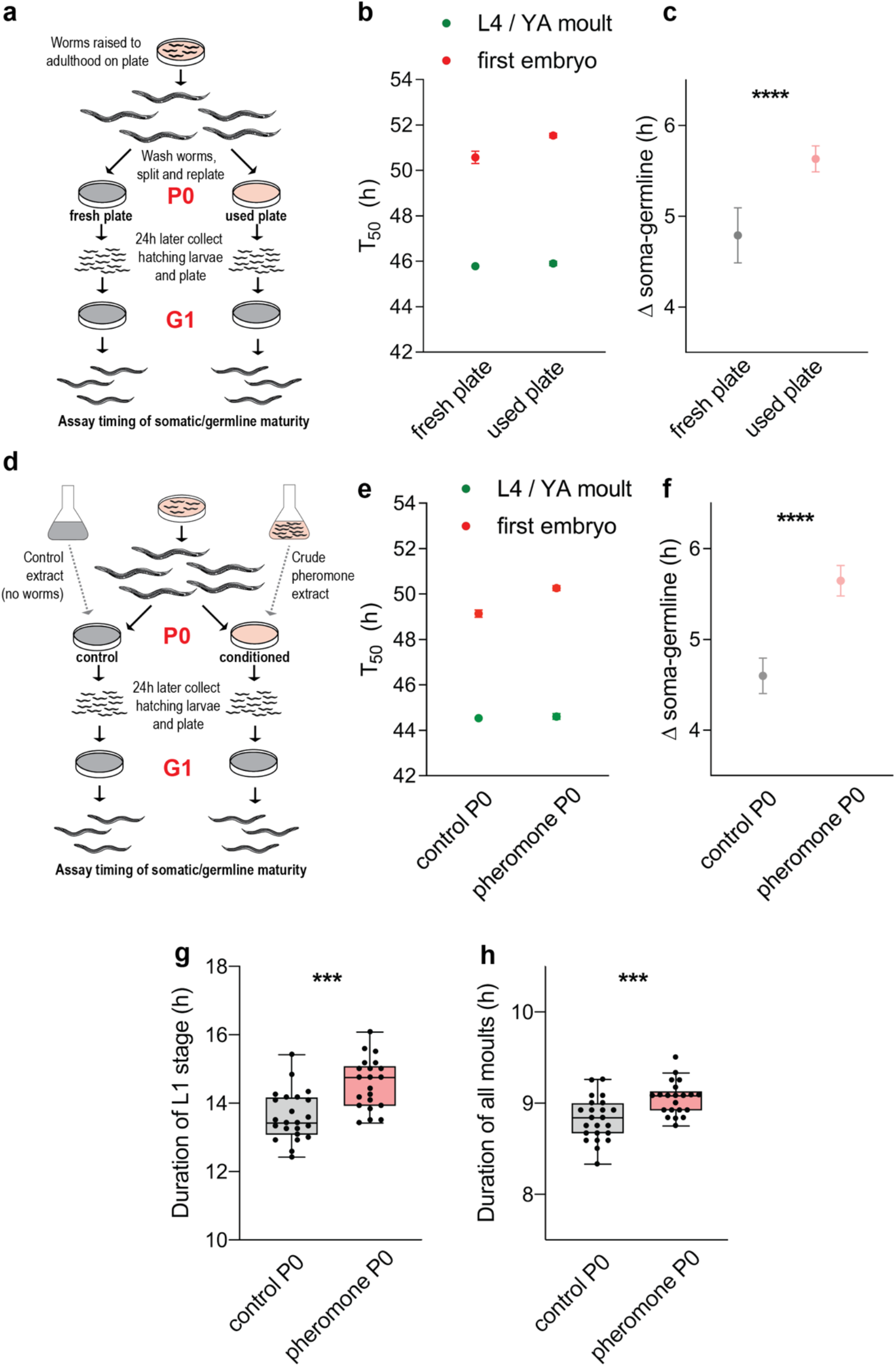
Parental exposure to media previously conditioned by worms leads to relatively delayed germline maturity and an increased brood size in progeny. **a)** Experimental schematic demonstrating the division of a synchronized adult population onto fresh plates or remaining on the used plate they were raised on, with progeny collected later for assaying developmental timings. **b)** Timings of the L4 to young adult molt (green), a proxy for somatic maturity, and the appearance of the first embryo (red), a proxy for germline maturity, for worms whose parents were moved to a fresh plate or remained on a used plate. T_50_ is the estimated time when half of the population passed through a developmental transition. T_50_ calculated based on counts of between193 and 346 worms (n = 193-346). **c)** The time between somatic maturity and germline maturity for worms whose parents were moved to a fresh plate or remained on a used plate, derived from the T_50_ measurements shown in panel **b**. This corresponds to the period of young adulthood where worms are adult but not yet gravid. **d)** Experimental schematic demonstrating the conditioning of plates with crude pheromone extract, produced by filtering liquid media in which worms had been cultured for 3-4 days. Parental worms were moved to plates conditioned with crude pheromone extract or control extract derived from a culture with no worms. **e)** Timings of the L4/YA molt (green) and first embryo (red) for worms whose parents were moved to plates conditioned with control extract or crude pheromone extract. n = 141-254. **f)** The time between somatic maturity and germline maturity for worms whose parents were moved to plates conditioned with control extract or crude pheromone extract, derived from the T_50_ measurements shown in panel **e**. **g**) Boxplot showing duration of the L1 larval stage for single worms whose parents were exposed to control (grey) or pheromone-conditioned (pink) plates. n= 23, 21. **h**) Boxplot showing the combined duration of the four moulting lethargus periods (M1-4) for single worms whose parents were exposed to control (grey) or pheromone-conditioned (pink) plates. n= 23, 21. All experiments were replicated at least 3 times independently. Panels **g** and **h** show data from a single representative replicate. Panels **b**, **c**, **e** and **f** show representative data from a single trial. Significance levels shown in panels **g** and **h** are from unpaired t tests on the data shown, whereas p values cited in main text are from linear mixed model analysis of 3 replicates pooled. Boxplots show Tukey whiskers with data points overlaid. Error bars in panels **b**, **c**, **e** and **f** show 95% confidence interval (note that they may not be visible when the error is small). P0, parental generation. G1, first progeny generation. YA, young adult. ***, p< 0.001 **\*\*\*\***, p < 0.0001.

This observation suggested the possibility that the parental environment influenced the phenotype of progeny. To investigate this further, we moved worms to fresh plates at various times prior to collection of progeny. We observed a maximum acceleration of germline maturity in progeny that hatched around 1 day after parents were transferred to fresh plates, after which the effect slowly decreased with the time the parents spend on the fresh plate (Fig S1a-d).

These observations led us to hypothesize that the effect of moving to a fresh plate vs remaining on a used plate is due to conditioning by pheromones secreted by worms: parents coming from used plates have been exposed to the pheromone accumulated since hatching, while parents moved to fresh plates are initially not exposed to pheromone, which then accumulates with time. However, these observations could also be explained by decreased food availability or quality caused by consumption. To exclude this possibility, we kept food supply constant across conditions by transferring adult worms onto fresh plates conditioned by either a bacteria-free crude pheromone extract from a dense worm liquid culture or by filtered bacteria-free medium from a bacterial culture as control. We collected a synchronized cohort of hatching progeny around 1 day later and immediately transferred it to fresh plates for development (Fig 1d). We found that parental exposure to crude pheromone extract delays the relative timing of germline maturation in progeny (Fig 1e, f), typically by 0.5-2 h. Although affected by maternal age (Perez et al. 2017), we found no reproducible impact of parental pheromone exposure on progeny brood size (p-value across 7 trials; 0.81; Fig S1e).

While there was a tendency for progeny of pheromone-treated worms to be delayed in reaching the adult molt, this varied substantially between trials on solid media. To clarify this, we used a previously published protocol that precisely measures somatic developmental timings in single worms in liquid culture (Olmedo et al. 2015). We found that progeny of pheromone-treated parents reached adulthood around 1 h later than progeny of control-treated parents (Fig S1f; Bonferroni-adjusted p-value across replicates = 7.06 × 10^−8^). This delay was largely due to an extension of the L1 larval stage by an average of 48 min (Fig 1g: adjusted p-value = 2.78 × 10^−13^). Other larval stages did not contribute significantly to this delay, either individually or in combination (Fig S1g; combined delay 1 min, adjusted p-value = 1 for all stages). Interestingly, however, around 25% of the total progeny developmental delay induced by exposure of parents to pheromone was due a slight extension of all 4 periods of molting lethargus, a sleep-like quiescent state (Raizen et al. 2008)(Fig 1h; adjusted p-value = 1.22 × 10^−5^). Taken together, these results suggest that the delay observed is largely due to a delay in developmental initiation rather than a slowing of developmental rate (Olmedo et al. 2020).

The phenotypes of the first progeny hatching on a new plate, laid shortly after parents were transferred, resembled those of worms hatched on the old plate, suggesting that it was an intergenerational effect of parental environment, rather than the environment encountered upon hatching, that determined the phenotype of progeny (Fig S1a-d). To confirm this, we performed two control experiments. In the first experiment, we harvested embryos laid by pheromone-exposed or control parents and briefly hypochlorite treated and washed them before allowing them to hatch on fresh plates (Fig S2a). In the second experiment, we harvested embryos laid by untreated parents and allowed them to hatch on either pheromone-conditioned or control plates (Fig S2d). In both cases it was the environment experienced by the parents, not the progeny upon hatching, which determined the delayed germline phenotype in progeny (Fig S2b, c, e-h).

### Exposure of parents to ascarosides is sufficient to induce germline delay in progeny

The observation that the extent of germline acceleration decreases with the time the parents spend on the new plate after ~ 1 day (Fig S1a-d) also suggests that the concentration of crude pheromone to which a hermaphrodite is exposed quantitatively modulates the extent of the germline delay phenotype observed in progeny. To test this, we performed a dose-response experiment that showed that indeed the extent of relative germline delay varied in an approximately log-linear fashion with pheromone concentration (Fig 2a, Fig S3a). This demonstrates quantitative control of progeny phenotypes in response to environmental cues perceived by parents.

**Figure 2.**
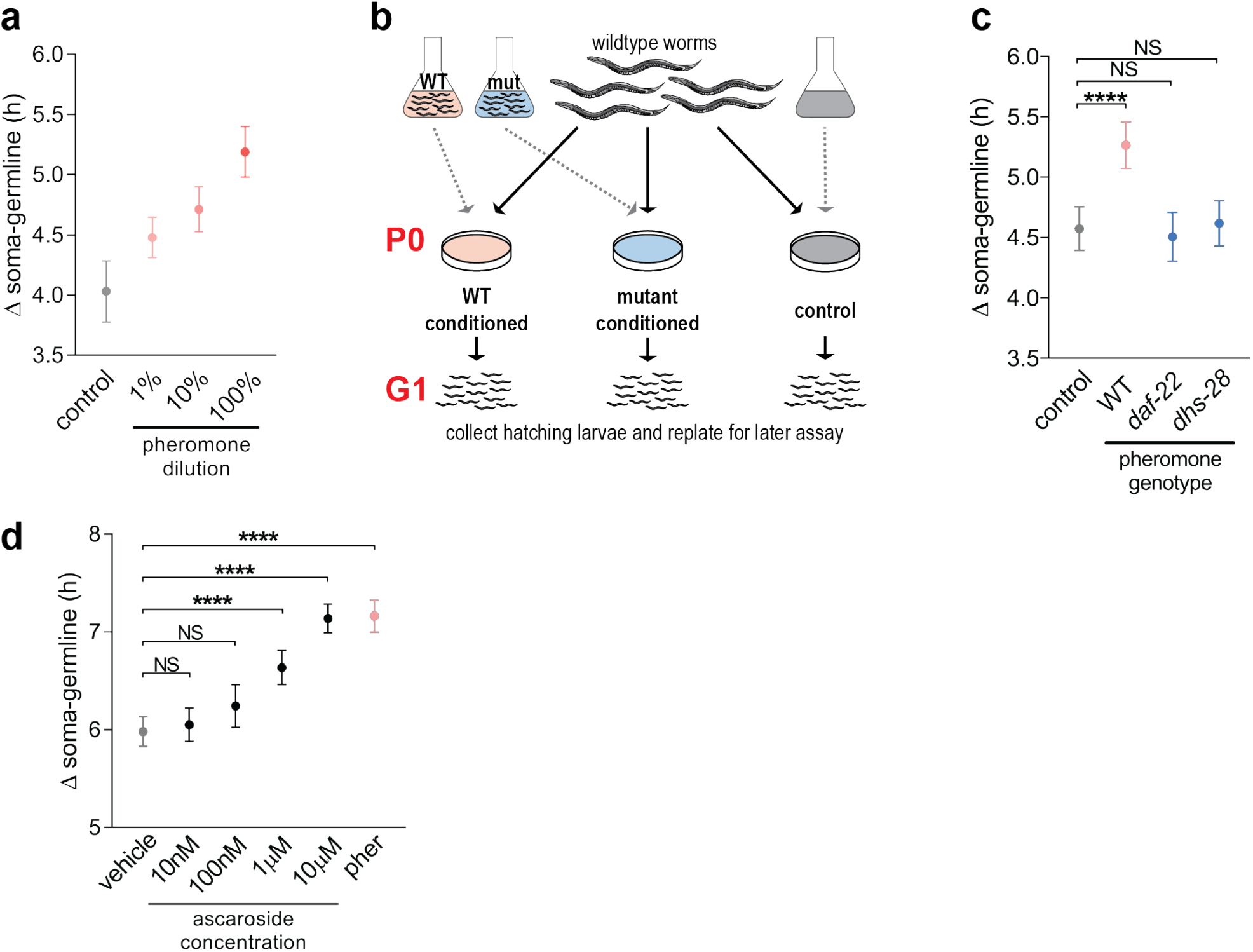
The effect of crude pheromone extract is dose-dependent and is likely to be mediated by major ascarosides. **a)** The time between somatic and germline maturity for worms whose parents were exposed to various dilutions of crude pheromone extract. L4/YA moltand first embryo T_50_ measurements are shown in **Fig S3a**. n = 214-429. **b)** Experimental schematic demonstrating the conditioning of plates with wildtype (pink) or mutant (blue) pheromone extract. Wildtype hermaphrodites are then exposed to control or conditioned plates and progeny are collected to assay developmental phenotypes. **c)** The time between somatic maturity and germline maturity for worms whose parents were exposed to control plates (grey), wildtype pheromone extract (pink), or *daf-22* or *dhs-28* mutant pheromone extract (blue). T_50_ measurements shown in **Fig S3b**. n = 137-283. **d)** The time between somatic maturity and germline maturity for worms whose parents were exposed to vehicle-only control plates (grey), various concentrations of an equimolar blend of synthetic major ascarosides #2 and #3, or crude pheromone extract. T_50_ measurements shown in **Fig S3c**. n = 191-337. All experiments were replicated at least 2 times independently. Panels **a**, **c** and **d** show representative data from a single trial. P0, parental generation. G1, first progeny generation. Pher, crude pheromone conditioned plates. NS, not significant. **\*\*\*\***, p < 0.0001. Error bars in panels **a**, **c** and **d**, 95% CI.

Which kind of pheromone is responsible for this intergenerational phenotype? The predominant pheromones characterized in *C. elegans* are the ascarosides, a group of small, secreted fatty acid-based molecules that mediate density perception upstream of dauer formation, among other functions (Ludewig and Schroeder 2018). To test the role of ascarosides we prepared crude pheromone extract from loss-of-function mutants for the metabolic genes *dhs-28* and *daf-22*, both required for the production of the biologically active short-chain ascarosides (Butcher et al. 2009; Fig 2b). Indeed, *dhs-28* and *daf-22* crude extracts are unable to elicit any germline delay in the progeny of exposed wildtype worms (Fig 2c; Fig S3b).

*daf-22* worms fail to produce hundreds of metabolites, not only ascarosides (Artyukhin et al. 2018). For this reason tested whether exposing parents to a synthetic blend of major ascarosides (#2,#3) would induce a relative germline delay in progeny. Indeed, we found that parental exposure to synthetic ascarosides was sufficient to elicit this intergenerational phenotype (Fig 2d; Fig S3c). Progeny response to parental ascaroside exposure was dose responsive in a log-linear fashion, with high concentrations of ascarosides eliciting a similar degree of relative germline delay to our crude pheromone extracts. Note however that we do not exclude any additional contribution by other *daf-22* or *dhs-28* dependent secreted metabolites.

### Pheromone perception in parental neurons mediates progeny phenotypes

Pheromones such as ascarosides are detected by chemosensory neurons. We tested if perception of pheromone is required in the parents to elicit the intergenerational effect of pheromone in the progeny. We mated hermaphrodites bearing loss-of-function mutations in neuronal-specific genes involved in pheromone signal transduction to wildtype males prior to pheromone exposure (Fig 3a). Control mothers are wildtype hermaphrodites mated to loss-of-function mutant males such that the genotypes of the assayed progeny of both groups were identical heterozygotes.

**Figure 3.**
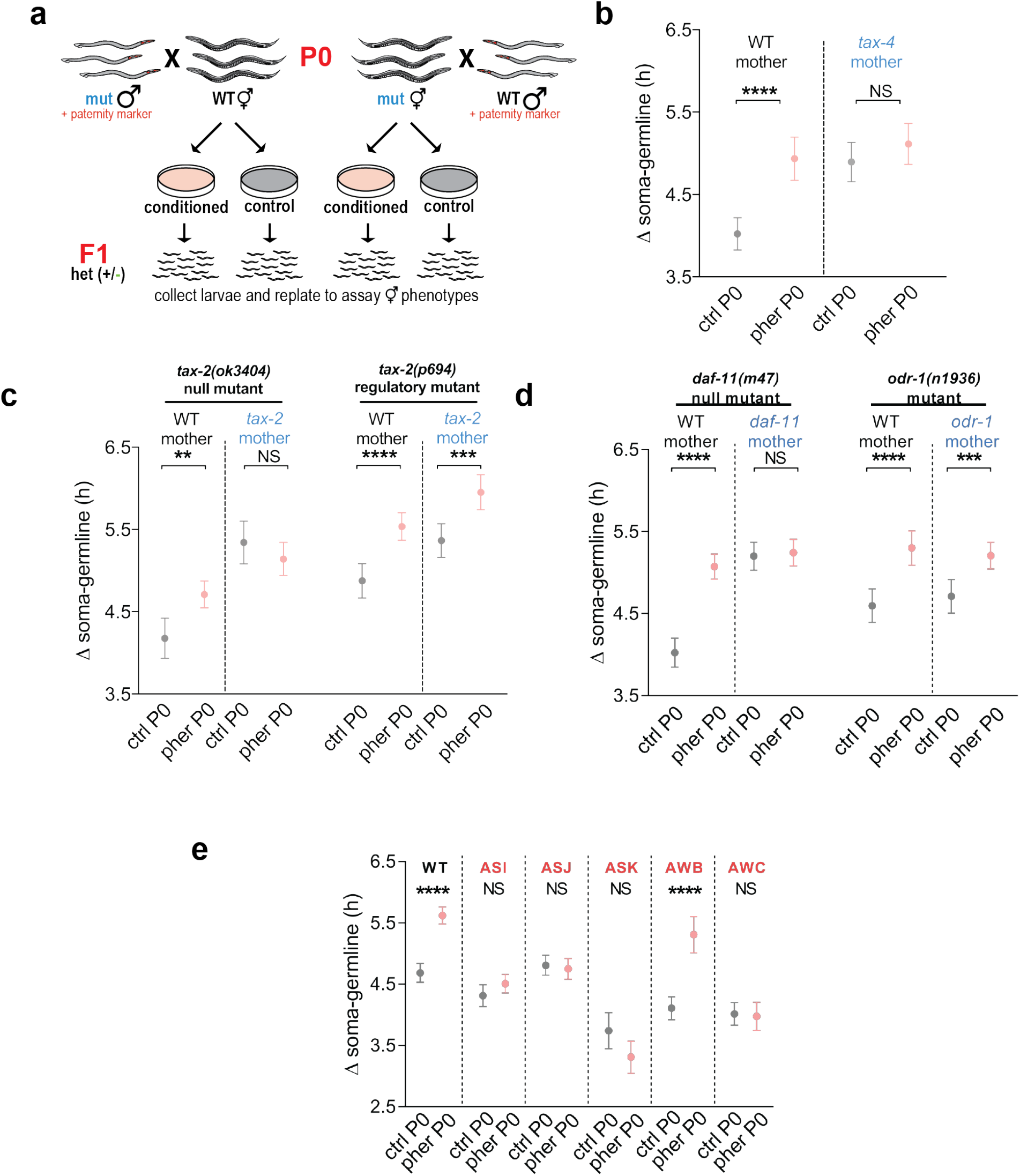
The TAX-2/TAX-4 cyclic nucleotide-gated channels and the upstream guanylyl cyclase DAF-11 are required in parental neurons for progeny response to exposure of parents to pheromone. **a)** Experimental schematic demonstrating crossing wildtype males to mutant hermaphrodites (and vice versa) prior to pheromone exposure. After crossing hermaphrodites are then exposed to control or conditioned plates and heterozygous progeny from crosses are collected to assay developmental phenotypes. Males carry a fluorescent paternity marker that allows confirmation of efficient crossing and removal of any self progeny. **b)** Time between somatic and germline maturity for *tax-4* heterozygous worms whose mothers were wildtype hermaphrodites (left) or homozygous *tax-4(ok3771)* mutant hermaphrodites (right) exposed to control plates (grey) or wildtype pheromone extract (pink). T_50_ measurements shown in **Fig S4a**. n = 81 - 210. **c)** Time between somatic and germline maturity for worms heterozygous for either the *tax-2(ok3404)* null allele (left half) or *tax-2(p694)* regulatory mutant (right half) whose mothers were wildtype hermaphrodites (left of dashed line) or homozygous *tax-4(ok3771)* mutant hermaphrodites (right of dashed line) exposed to control plates (grey) or wildtype pheromone extract (pink). T_50_ measurements shown in **Fig S4b**. n = 129-250. **d)** Time between somatic and germline maturity for worms heterozygous for mutant alleles of the receptor-bound guanylyl cyclases *daf-11(m47)* (left half) or *odr-1(n136)* (right half) whose mothers were wildtype hermaphrodites (left of dashed line) or homozygous mutant hermaphrodites (right of dashed line) exposed to control plates (grey) or wildtype pheromone extract (pink). T_50_ measurements shown in **Fig S4c**. n = 57-271. **e)** Time between somatic and germline maturity for worms with genetic ablations of the ASI, ASJ, ASK, AWB or AWC chemosensory neurons exposed to control plates (grey) or wildtype pheromone extract (pink). T_50_ measurements shown in **Fig S4d**. n = 125-264. All experiments were replicated at least 2 times independently. All panels show representative data from a single trial. All data shown in a single panel are from the same trial. P0, parental generation. F1, first filial generation. WT, wildtype. Ctrl, control-conditioned plates. Pher, crude pheromone conditioned plates. NS, not significant. **\*\***, p < 0.01. **\*\*\***, p < 0.001. **\*\*\*\***, p < 0.0001. Pairwise tests are subject to Bonferroni corrections for multiple testing within each trial. Error bars in panels **b**, **c**, **d** and **e** 95% CI.

We first tested *tax-2* and *tax-4*, encoding two subunits of a cyclic nucleotide-gated ion channel previously implicated in signal transduction downstream of pheromone in chemosensory neurons (Bargmann 2006). We found that progeny of *tax-2* and *tax-4* mutants already displayed delayed germline maturation relative to wildtype under control conditions, and indeed do not respond with a further delay to parental pheromone exposure (Fig 3b, c; Fig S4a, b). We went on to test mutants of two guanylate cyclases, DAF-11 and ODR-1, that relay signals from the GPCRs that detect environmental stimuli to control TAX-2/TAX-4 ion channel function (Bargmann 2006). We found that the progeny of *daf-11* parents phenocopy the progeny of *tax-2/tax-4* parents, showing a constitutive germline delay that failed to respond further to parental pheromone exposure (Fig 3d; Fig S4c). This was in keeping with the reported role of DAF-11 in counteracting dauer formation downstream of pheromone exposure (Birnby et al. 2000). In contrast, progeny of mutants for *odr-1*, which does not affect dauer formation (Noelle and Bargmann 2000), did respond to parental pheromone exposure (Fig 3d; Fig S4c).

To identify which neurons were required for the intergenerational response to pheromone, we tested the *tax-2* regulatory mutant *p694*, which compromises TAX-2 function in a subset of chemosensory neurons (Coburn and Bargmann 1996). We found that this mutant displayed a robust intergenerational response to pheromone (Fig 3c; Fig S4b). Of the 6 pairs of chemosensory neurons in which *tax-2* functions in the *p694* mutant, 5 pairs also express *daf-11* (Birnby et al. 2000): the ASI, ASJ, ASK, AWB and AWC neurons. We tested genetic ablations of these 5 specific pairs of neurons. The results show that the ASK and ASI neurons are both required for the intergenerational germline delay response to pheromone (Fig 3E, Fig S4d). The AWB neurons were not required for germline delay. The germline delay was largely abrogated or absent for the ASJ and AWC ablation strains in most trials, but we note that both did show a significant pheromone-induced delay in 1/5 and 1/4 trials respectively.

### Germline delay results from delayed initiation of post-embryonic development

What is the developmental origin of the soma-germline heterochrony that we observe at adulthood? A recent study found that the developmental delay arising after recovery from starvation-induced L1 arrest occurred not due to a slowing of developmental rate but rather due to a delay in initiation of post-embryonic development upon feeding (Olmedo et al. 2020). This led us to hypothesize that pheromone exposure in the parents might impact the relative timing of post-embryonic developmental initiation in progeny somatic or germline tissues, rather than their relative rate of development. Further, this hypothesis could explain the intergenerational germline delay as a latent effect of maternal control over an early developmental event, whose effects manifest in progeny up to adulthood. To test this hypothesis, we monitored the timings of initial postembryonic divisions of the primordial germ cells (PGCs) Z2 and Z3, the M mesoblast cell and the head (H) lineage of the hypodermal seam cells (Fig 4a).

**Figure 4.**
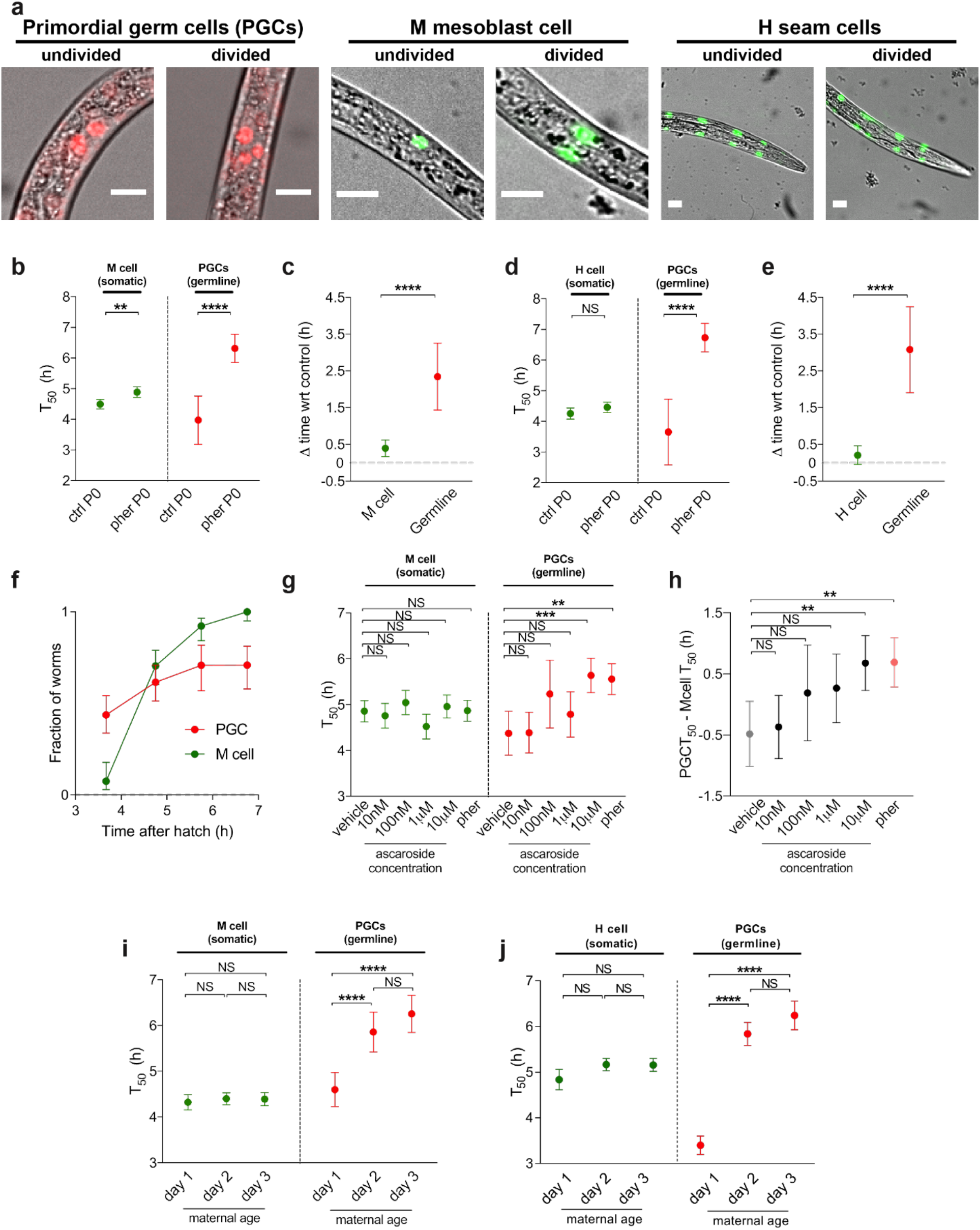
Initiation of germline development is delayed in progeny of worms exposed to pheromone. **a)** Microscopy images showing the primordial germ cells (PGCs), somatic M cell (mesoblast precursor) and somatic H cells (hypodermal seam cell precursor) before and after initial divisions during L1 stage larval development. Merged images show brightfield and mCherry fluorescence in red (PGCs) or GFP fluorescence in green (M cell and H seam cells). Scale bar for all images 10 μm. **b)** Timings of initial divisions of the somatic M cell (green, left of dashed line) and PGCs (red, right of dashed line) for progeny of parents exposed to control or pheromone conditioned plates. n = 48-109. **c)** Impact of pheromone on timings of initial division of the somatic M cell (green) or PGCs (red). Impacts are calculated from data shown in panel **b**. **d)** Timings of initial divisions of the somatic H cells (green, left of dashed line) and PGCs (red, right of dashed line) for progeny of parents exposed to control or pheromone conditioned plates. n = 51-98. **e)** Impact of pheromone on timings of initial division of the somatic H cells (green) or PGCs (red). Impacts are calculated from data shown in panel **d**. **f)** Raw counts of individuals with divided M cells (green) or PGCs (red) at various time points for data shown in panel **b**. **g)** Timings of initial divisions of the somatic M cell (green, left of dashed line) and PGCs (red, right of dashed line) for progeny of parents exposed to control plates, increasing concentrations of synthetic ascarosides #2 and #3 or crude pheromone extract. n = 87-224. **h)** The time between T_50_s for the PGCs and the somatic M cell reporter for progeny of parents exposed to control plates (grey), increasing concentrations of synthetic ascarosides #2 and #3 (black) or crude pheromone extract (pink), calculated from data from panel **g**. **i)** Timings of initial divisions of the somatic M cell (green, left of dashed line) and PGCs (red, right of dashed line) for progeny of mothers on day 1, 2 or 3 of adulthood. n= 93-253. **j)** Timings of initial divisions of the somatic H cell (green, left of dashed line) and PGCs (red, right of dashed line) for progeny of mothers on day 1, 2 or 3 of adulthood. n = 63-141. All experiments were replicated at least 2 times independently. All panels show representative data from a single trial. All data shown in a single panel are from the same trial. PGCs, primordial germ cells. Ctrl, control-conditioned plates. Pher, crude pheromone conditioned plates. P0, parental generation. Wrt, with respect to. NS, not significant. **\*\***, p < 0.01. **\*\*\***, p < 0.001. **\*\*\*\***, p < 0.0001. Pairwise tests are subject to Bonferroni corrections for multiple testing within each trial.

We observed that while parental exposure to pheromone had a modest impact on the timings of somatic M cell division and a negligible impact on H seam cell division, the initial division of the PGCs is strongly delayed (Fig 4, b-e). Interestingly, we observed that inter-individual variability within groups was much more pronounced for initial PGC divisions than for somatic cell divisions (Fig 4f). Importantly, parental exposure to a blend of synthetic ascarosides #2 and #3 led to a dose-responsive slowing of PGC division relative to M cell division (Fig 4g, h), with the highest concentrations having a similar impact to crude pheromone extract.

### Pheromone perception is required for maternal age-dependent changes in progeny heterochrony

We previously reported maternal age as influencing soma-germline heterochrony in adulthood (Perez et al. 2017). Can this be explained by differential timings of initial post-embryonic divisions in somatic and germline tissues? Indeed, maternal age had a negligible effect on the timing of the initial division of the M cell and H cells but had a strong effect on the division of the PGCs, which were delayed in progeny of older mothers (Fig 4i, j). While this observation can explain the relative germline delay with respect to somatic maturity observed in adulthood in progeny of older mothers, these worms still tend to reach somatic and germline maturity sooner than early progeny (Perez et al. 2017), implying an overall acceleration of development.

We also previously reported that while many phenotypic effects of maternal age could be explained by differential provisioning of the yolk lipoprotein complex vitellogenin, it was unlikely to be the cause of the observed soma-germline heterochrony (Perez et al. 2017). Further, parental vitellogenin provisioning to embryos is not altered in response to pheromone exposure (Fig S5a). Therefore, we wondered whether the apparent effect of maternal age on soma-germline heterochrony could be explained not by age *per se* but by elevated pheromone exposure in older mothers. To test this hypothesis, we assayed soma-germline heterochrony in adulthood in *tax-4* mutants, in which pheromone is unable to induce intergenerational germline delay. First, we showed that *tax-4* worms were capable of conditioning media to induce intergenerational germline delay in wildtype worms (Fig S5b, c). Further, we confirmed by examining early embryo autofluorescence (Perez et al. 2017) that *tax-4* worms displayed an increase in embryonic vitellogenin loading with maternal age (Fig S5d). Likewise, progeny length at hatch, previously shown to be vitellogenin-dependent, increased with maternal age in *tax-4* mutant worms (Fig S5e). However, *tax-4* mutants did not display a delayed germline in the progeny of the oldest mothers (Fig S5f, g). This indicates that the previously-reported alteration of progeny soma-germline heterochrony with maternal age is indeed pheromone-dependent, rather than an intrinsic consequence of maternal age.

### Inheritance requires DAF-7/TGF-β in mothers and DAF-12/NHR and DAF-3/Co-SMAD in progeny

To better understand how the effects of parental pheromone exposure are inherited, we tested whether progeny sired by pheromone-exposed males also exhibit a germline delay (Fig 5a). We used sperm-less *fog-2* mutant females to ensure that all progeny were sired by males, and wildtype males so that assayed progeny were *fog-2(+/-)* hermaphrodites that were able to self-fertilize. We found no evidence for transmission of paternal pheromone exposure via sperm in male-sired progeny (Fig 5b; Fig S6a). The converse experiment, with females exposed to pheromone, confirmed that memory of maternal pheromone exposure is transmitted through the oocyte (Fig 5b; Fig S6a). However, we cannot exclude inheritance via sperm, due to two caveats. First, we did not test transmission via sperm in selfing hermaphrodites, due to the difficulty of cleanly distinguishing maternal and paternal transmission. Second, both pheromone production and responses are sex-specific in *C. elegans* (Chute and Srinivasan 2014), meaning a different physiological response to pheromone in males could also underlie a failure to observe transmission to their progeny.

**Figure 5.**
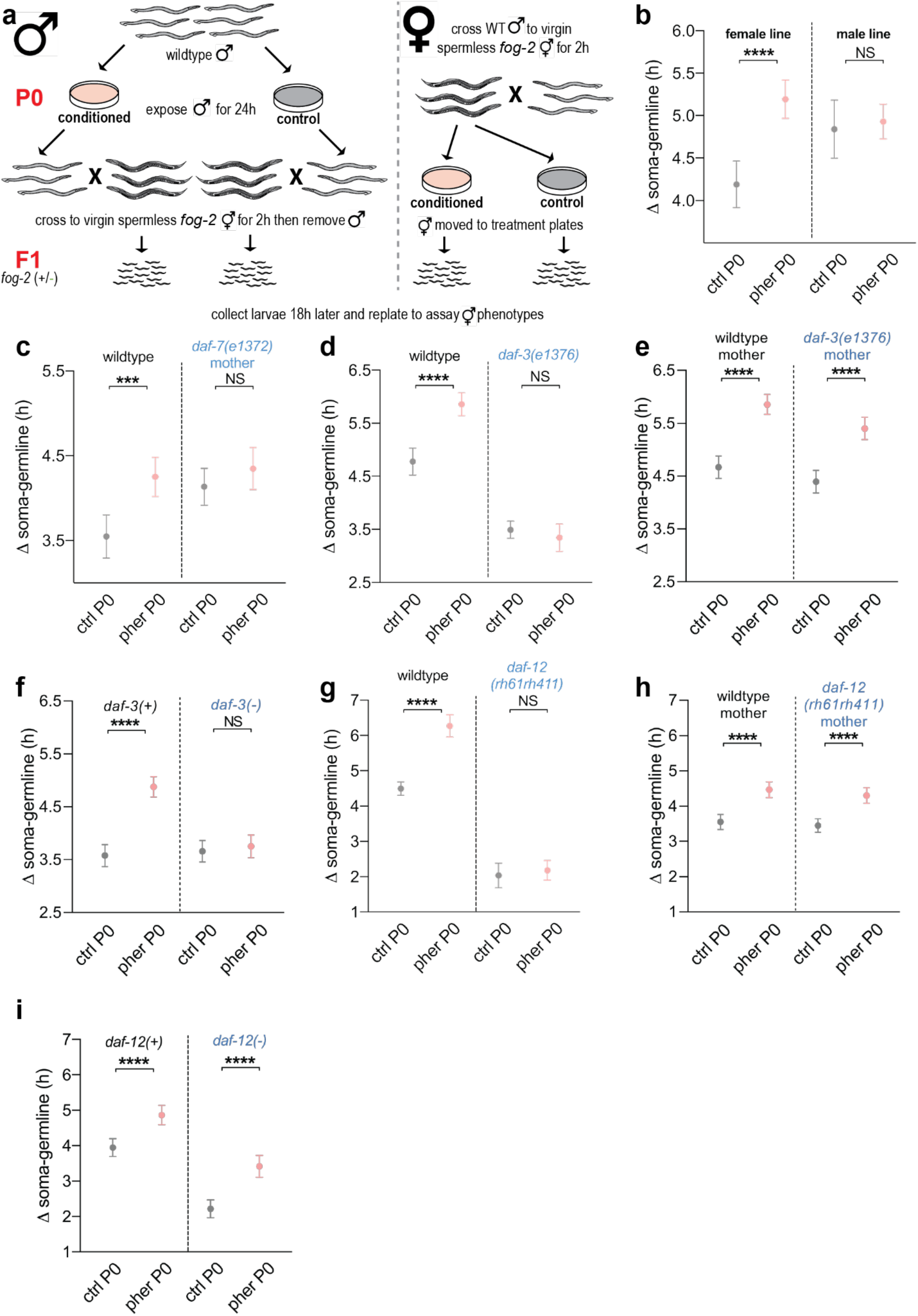
Effects of parental pheromone exposure are inherited via the oocyte and require DAF-7/TGF-β in parental neurons and the Co-SMAD DAF-3 and nuclear hormone receptor DAF-12 in progeny. **a)** Schematic showing experiment to test for transmission of parental pheromone exposure via male sperm (left of dashed line). The converse experiment, to confirm transmission via the oocyte, is also shown (right of dashed line). **b)** Time between somatic and germline maturity for male-sired hermaphrodites whose mothers (left of dashed line) or fathers (right of dashed line) were exposed to control plates (grey) or crude pheromone (pink). T_50_ measurements shown in **Fig S6a**. n = 94-154. **c)** Time between somatic and germline maturity for male-sired wildtype worms (left of dashed line) or male-sired heterozygous progeny of *daf-7(e1372)* mutant mothers (right of dashed line) whose mothers were exposed to control plates (grey) or crude pheromone (pink). T_50_ measurements shown in **Fig S6b**. n = 102-200. **d)** Time between somatic and germline maturity for wildtype worms (left of dashed line) or *daf-3(e1376)* mutant worms (right of dashed line) whose parents were exposed to control plates (grey) or crude pheromone (pink). T_50_ measurements shown in **Fig S6c**. n = 120-242. **e)** Time between somatic and germline maturity for *daf-3* heterozygous progeny whose wildtype mothers (left of dashed line) or *daf-3(e1376)* mutant mothers (right of dashed line) were exposed to control plates (grey) or crude pheromone (pink). T_50_ measurements shown in **Fig S6d**. n = 73-233. **f)** Time between somatic and germline maturity for *daf-3(+)* (left of dashed line) or homozygous *daf-3(e1376)* mutant (right of dashed line) progeny whose *daf-3* heterozygous mothers were exposed to control plates (grey) or crude pheromone (pink). T_50_ measurements shown in **Fig S6e**. n = 85-135. **g)** Time between somatic and germline maturity for wildtype worms (left of dashed line) or *daf-12(rh61rh411)* mutant worms (right of dashed line) whose parents were exposed to control plates (grey) or crude pheromone (pink). T_50_ measurements shown in **Fig S6f**. n = 102-208. **h)** Time between somatic and germline maturity for *daf-12* heterozygous progeny whose wildtype mothers (left of dashed line) or *daf-12(rh61rh411)* mutant mothers (right of dashed line) were exposed to control plates (grey) or crude pheromone (pink). T_50_ measurements shown in **Fig S6g**. n = 63-181. **i)** Time between somatic and germline maturity for *daf-12(+)* (left of dashed line) or homozygous *daf-12(rh61rh411)* mutant (right of dashed line) progeny whose *daf-12* heterozygous mothers were exposed to control plates (grey) or crude pheromone (pink). T_50_ measurements shown in **Fig S6h**. n = 67-194. All experiments were replicated at least 2 times independently. All panels show representative data from a single trial. All data within a panel comes from the same trial. Error bars in panels show 95% confidence interval. Pairwise tests are subject to Bonferroni corrections for multiple testing within each trial. Ctrl, control-conditioned plates. Pher, crude pheromone conditioned plates. P0, parental generation. F1, first filial generation. NS, not significant. **\*\*\***, p < 0.001. **\*\*\*\***, p < 0.0001.

High pheromone concentrations can suppress germline proliferation in adult hermaphrodites by downregulating the expression of the TGF-β ligand *daf-7* in ASI neurons (Dalfó et al. 2012). Additionally, *daf-7* expression has been shown to be downstream of *daf-11*, *tax-2* and *tax-4* in multiple contexts (Meisel et al. 2014, Murakami et al. 2001) and has been implicated in transgenerational inheritance (Kaletsky et al. 2019, Moore et al. 2019). We wondered whether this established neuron-to-germline signaling axis might also contribute to the intergenerational inheritance that we observe. As *daf-7* mutants are dauer-constitutive, we crossed wildtype males to *daf-7(e1372)* mutant hermaphrodites (Ren et al. 1996) to ensure wildtype-like development in heterozygous progeny before pheromone exposure. Note that as *daf-7* mutant males are defective for crossing, the controls assayed in this experiment are male-sired homozygous wildtype worms, rather than *daf-7* heterozygotes. Like *tax-2/4* and *daf-11* mutants, progeny of *daf-7* mutant mothers have a constitutively delayed germline, consistent with low *daf-7* expression signaling high environmental pheromone concentration, and they do not respond further to maternal pheromone exposure (Fig 5c; Fig S6b).

The principal downstream effector of the *daf-7* TGF-β pathway is DAF-3, a Co-SMAD whose activity is antagonized by DAF-7 (Gumienny and Savage-Dunn 2018). As such, *daf-3* mutants phenocopy high *daf-7* expression and display superficially wildtype development in benign conditions, with males competent for crossing. We exposed *daf-3(e1376)* null mutants to pheromone- or control-conditioned plates and assayed their progeny. The germline in *daf-3* mutants is constitutively advanced with respect to the soma (Fig 5d). This appears to result primarily from a delay in somatic, but not germline, development (Fig S6c). Like *daf-7* mutants, however, *daf-3* mutants do not display any disproportionate germline delay in response to parental pheromone exposure (Fig 5d). We hypothesized that DAF-3 might act in the maternal somatic gonad downstream of DAF-7 to influence the germline and the phenotypes of the next generation (Pekar et al. 2017). However, rescuing *daf-3* function in progeny restored pheromone-induced germline delay in the progeny of mutant mothers (Fig 5e; Fig S6d), indicating that DAF-3 is dispensable in parents and acts in progeny to facilitate intergenerational germline delay. Indeed, *daf-3*(-/-) progeny of *daf-3* heterozygous mothers do not display a germline delay in response to parental pheromone exposure, unlike their *daf-3(+/-)* siblings (Fig 5f, Fig S6e).

Another factor that responds to pheromone downstream of DAF-7/TGF-β, and possibly DAF-3 (Jia et al. 2002), in the dauer fate decision is the nuclear hormone receptor DAF-12. *daf-12* mutants display extensive heterochronic phenotypes during development (Antebi et al. 2000) and *daf-12* has been proposed to contribute to developmental robustness (Hochbaum et al. 2011). *daf-12(rh61rh411)* null mutants do not display germline delay in response to parental pheromone exposure (Fig 5g). *daf-12* mutants have an extremely advanced germline relative to somatic development (Fig 5g), with embryos appearing shortly after the completion of L4/YA molt. Like *daf-3* mutants, this appears to be a result of the mutation introducing a substantial delay in the completion of somatic, but not germline, development (Fig S6f).

We rescued *daf-12* function in progeny and found that DAF-12 is not necessary in parents for pheromone-induced intergenerational germline delay (Fig 5h; Fig S6g). However, wildtype *daf-12* in heterozygous mothers can rescue the pheromone-induced germline delay in *daf-12* (-/-) mutant progeny (Fig 5i; Fig S6h). However, although maternally-rescued *daf-12*(-/-) worms are responsive to parental pheromone perception, they still display a constitutively advanced germline (Fig 5i; Fig S6h). From this we conclude that DAF-12 controls soma/germline relative timings within the same generation, as well as being necessary for the pheromone-induced intergenerational germline delay. The later effect, though not the former, can be rescued by a maternal copy of *daf-12*.

Most upstream pathways that converge on DAF-12 act through the cytochrome P450 DAF-9, which produces small molecule DAF-12 ligands known as dafachronic acids (Motola et al. 2006). We hypothesized that dafachronic acid itself could be inherited through the oocyte in order to influence progeny developmental initiation through modulating the activity of DAF-12. To test this, we used *daf-9(m540)* loss-of-function mutant hermaphrodites (Jia et al. 2002). These worms were raised to adulthood on plates supplemented with (25S)-Δ7-dafachronic acid (DA) to bypass the constitutive dauer phenotype and induce wildtype-like development. We crossed these *daf-9* mutant hermaphrodites to wildtype males to ensure that the resulting *daf-9(+/-)* progeny also displayed wildtype-like development. We then moved the *daf-9* mutant mothers to plates supplemented with 0 nM (vehicle only), 10 nM or 100 nM DA, or to pheromone- or control-conditioned plates with no DA and collected their progeny to assay phenotypes. We found that parental DA feeding had no effect on progeny somatic or germline development (Fig S6i, j). Indeed, exposure of *daf-9* mutant mothers to pheromone induced a robust germline delay in their progeny (Fig S6i, j), invalidating the hypothesis that DA is the inherited signal.

The lack of requirement in parents for either DAF-3 or DAF-9/DAF-12 suggests that parental DAF-7 controls progeny germline development via a non-canonical signaling axis.

## DISCUSSION

We have shown here that neuronal perception of the environment can intergenerationally control the development and generation time of an animal. Specifically, neuronal sensing of a social cue transmits a signal that crosses the Weismann barrier to the germline to control the post-embryonic initiation of germline development in the next generation. Inheritance via the oocyte required the TGF-β ligand DAF-7 in parents and its targets DAF-3/CoSMAD and DAF-12/NHR in progeny (Fig S7a). Neuronal perception of pheromone also underlies the previously reported control of soma-germline heterochrony by maternal age, meaning that maternal age determines phenotypic variation intergenerationally by at least three signals in this species (Fig S7b).

The pheromone-induced intergenerational germline delay that we describe is likely to have a considerable impact on fitness. Minimum generation time is a critical fitness parameter in *C. elegans*. Indeed, a mutant that displayed a 50% increase in total fecundity at the cost of a modest extension of minimum generation time of 2-3 h was consistently outcompeted by wildtype worms in an ‘eating race’ (Hodgkin and Barnes 1991). *C. elegans* undergo rapid population expansion, rarely lasting for more than 3 or 4 generations (Félix and Duveau 2012), on ephemeral resource patches, followed by widespread entry into the dauer diapause awaiting the dispersal of a few individuals to a new resource patch (Frézal and Félix 2015). For population founders, minimizing the generation time at any cost to take advantage of plentiful resources during the exponential phase of population expansion is likely to be a high-fitness strategy. As the resource patch nears depletion, this strategy may no longer be optimal. Pheromone is likely to be a vital cue for worms looking to adapt their life history traits, and that of their progeny, to the appropriate stage of the population cycle.

Previous work in the nematode *Pristionchus pacificus* has shown that pheromone can indicate the age structure of a burgeoning population to adaptively influence progeny developmental outcomes. The composition of whole pheromone in this species is stage-specific, such that only pheromone produced by adults, not juveniles, can elicit an alternative developmental trajectory, a predation-enabling mouth morphology, in developing worms (Werner et al. 2018). In contrast, our study demonstrates direct inheritance of parental pheromone perception to alter progeny phenotypes. Given the importance of conspecific competition to organisms of all kinds, we speculate that such mechanisms to alter progeny phenotypes in response to the size and state of the population are widespread.

In summary, we have shown intergenerational control of the developmental timing of a specific tissue in an animal, in response to neuronally-perceived environmental cues experienced by the previous generation. This example, which controls the generation time of the animal, adds to the growing body of evidence (Baugh and Day 2020, Dias and Ressler 2014, Gapp et al. 2014a, Gapp et al. 2014b, Remy 2010, Burton et al. 2017) that neuronal perception of the environment can be inherited across generations and be an important influence on animal phenotypes and fitness.

## METHODS

### Worm culture

All worms were cultured using standard methods at 20°C. Worms were fed on nematode growth medium (NGM) plates seeded with *Escherichia coli* OP50. Worm strains used in this study are listed in Supplemental Table 1. Some strains were provided by the CGC, which is funded by NIH Office of Research Infrastructure Programs (P40 OD010440). The ASJ ablation strain ZD763 was a kind gift of Dennis Kim. Day 1, 2 and 3 mothers were synchronized as in (Perez et al. 2017). Mothers of the *tax-4* mutant strain VC3113 were approximately 12 h older than their wildtype counterparts, due to a developmental delay of a similar extent.

### Staging somatic and germline development in vivo

Molting and appearance of the first fertilized egg were used as developmental transitions to stage somatic and germline development, respectively, as in (Perez et al. 2017). Briefly, transitions were monitored in animals on agar plates using brightfield microscopy with an MZ9S stereo microscope (Leica). The fraction of the population that had undergone a developmental transition was estimated by counting worms before or after the transition, at timepoints before and after 50% of the population had undergone the transition. This count data was then modelled using a binomial generalized linear model, and the time at which half of the population had undergone the transition (T_50_) was estimated using the function ‘dose.p’ of the library ‘MASS’ in R. 95% confidence intervals were estimated by multiplying the standard error obtained by the ‘dose.p’ function by ±1.96. Δ soma-germline values for each group were calculated by subtracting the germline T_50_ from the soma T_50_. 95% confidence intervals for the Δ soma-germline values were estimated by first calculating a standard error by taking the square root of the sum of the two squared T_50_ standard errors, and then multiplying by ±1.96. The significance of the difference between two calculated T_50_ or Δ soma-germline values was determined by using the ‘comped’ function in the library ‘drc’ in R, as in (Perez et al. 2017). Bonferroni corrections were used to adjust significance thresholds to allow for multiple comparisons within a trial.

### Crude pheromone preparation

Crude pheromone was prepared by growing worms in large-scale liquid culture in liquid NGM medium. Worms from a small starting liquid culture were bleached and the embryos allowed to hatch overnight in M9 to arrest development at L1. Worms were then seeded into a 250 ml liquid culture in a 2.5 L Erlenmeyer flask at a density of 5-10 worms per μl. Streptomycin-resistant OP50-1 food that had previously been grown overnight in LB medium before washing and resuspension at high density was added to a final concentration of 50 mg ml^−1^, in the presence of 25 μg ml^−1^ streptomycin. Worms were grown at 20 °C with shaking at 180 rpm with additional food being added as necessary to avoid starvation for 4 days before harvesting pheromone, resulting in a population of gravid adults and abundant hatched larvae. Control cultures were grown in parallel with OP50-1 in liquid NGM supplemented with streptomycin but without any worms added. To harvest, cultures were spun gently at 200 *g* for 1 min several times to remove worms from suspension without any worm lysis. The cloudy supernatant was then spun for 15 minutes at 3000 *g* to pellet remaining bacteria. The clear supernatant was then passed through a 0.22 μm filter to exclude any remaining bacteria to yield the final crude pheromone preparation. This preparation was frozen and stored in 1 ml aliquots at −70 °C. For the bulk of the experiments, 3 large pheromone preparations were produced independently to use as biological replicates. Where smaller pheromone preparations were produced (e.g. in the case of *dhs-28* or *daf-22* mutant pheromone), a small batch of wildtype pheromone was produced in parallel for use as control in the relevant experiments.

### Conditioned plate assay

To set up a synchronized parental generation, around 20 synchronized gravid day 1 adults were allowed to lay eggs on a 90 mm NGM plate for 2 h before being removed. Approximately 65 h later, when these eggs have developed into young adult or newly gravid worms, they were washed with M9 buffer into a plastic 15 ml tube and spun down at 200 *g* before the supernatant was aspirated to 100 μl. Worms were then distributed drop by drop with a pipette tip rinsed with 0.1% Triton-X (Sigma Aldrich) between a new, fresh plate and the original, conditioned plate. Typical adult densities ranged from 50-200 worms per plate. Plates were allowed to dry before worms were incubated at 20 °C. 18-26 h later, plates were washed several times with M9 medium to remove adults and any hatched larvae, which were discarded. Embryos remaining on the plates were allowed to hatch for 1-2 h before newly hatched larvae were collected in M9 buffer, spun down at 200 *g* and transferred to fresh 90 mm plates for development. As these worms hatched on a plate with food they were never starved at any point. Typical larval densities ranged from 100-500 worms per plate. Both parental and larval densities varied between trials but were kept constant within trials. Care was taken to immediately remove any adults or embryos which may have contaminated the preparation of hatching worms. Larger contaminating larvae could be easily removed at a later point prior to assay. Somatic and germline development were assayed as described above.

Experiments with crude pheromone were performed as described above, substituting control- or pheromone-treated plates for new and conditioned plates. To prepare pheromone or control plates, 300 μl of a freshly thawed pheromone aliquot was applied to the lawn of a 60 mm plate by pipetting, and spread to cover the entire surface of the plate by tilting and rotating the plate. The plate was allowed to dry and then used within 1 h after application of pheromone.

Experiments with synthetic ascarosides #2 and #3 were also performed as above. Synthetic ascarosides were a kind gift of Frank Schroeder. Asc#2 and asc#3 stock solutions were 5 μM in 100% ethanol. Plates were prepared with an equimolar mix of asc#2 and asc#3, which was added to 300 μl of control supernatant and applied to plates as above. The ascaroside concentration reported in the figures represents the molarity of both ascarosides combined e.g. 10 mM refers to 5 mM each of asc#2 and asc#3. The ethanol concentration of control and diluted ascaroside solutions were adjusted by addition prior to application to plates so that ethanol concentration was held constant across treatments.

### Early life exposure control experiments

We performed control experiments to confirm that parental pheromone exposure, rather than early-life exposure of progeny to pheromone, was responsible for the observed phenotypes. In one control experiment (as shown in the schematic in Fig S2a), we prepared larvae that had hatched in the absence of pheromone, although their parents had been treated on pheromone or control plates. Embryos laid on plates by pheromone- or control-treated parents were scraped gently from the surface of the plate with a glass coverslip and collected after prior removal of adults and larvae. A mild alkaline hypochlorite treatment (‘bleaching’) was applied to kill any remaining hatched larvae before 3 washes with M9 buffer. Embryos thus prepared were then transferred onto a fresh plate. At least 2 h later, newly hatched larvae were removed, and embryos were allowed to hatch for a further 1-2 h. Larvae hatched in this time were collected and transferred to fresh plates for development and assay as described for the conditioned plate assay.

In a second control experiment (as shown in the schematic in Fig S2d), we prepared larvae that had hatched in the presence or absence of pheromone, although they were derived from the same untreated parental cohort. Parental worms were not transferred to pheromone or control plates but remained on the plate on which they had developed. Embryos were then scraped and briefly bleached as above. These embryos were then transferred to recently prepared pheromone and control plates. At least 2 h later, newly hatched larvae were removed, and embryos were allowed to hatch for a further 1-2 h. Larvae hatched in this time were collected and transferred to fresh plates for development and assay as above.

### Crossing experiments to assay mutation effect in the parental generation

In several assays, mutant hermaphrodites later treated with pheromone or control plates were first crossed to wildtype males prior to treatment, such that assayed progeny would be heterozygous for the mutation (as shown in the schematic in Fig 3a). This was to more confidently ascribe the intergenerational effects of the mutant gene to its action in the parental generation, as the assayed progeny would carry one functional wildtype allele, as well as in order to avoid mutant phenotypes which complicated assaying homozygous mutant progeny, such as slow or heterogeneous rates of development or widespread dauer entry (such as in *daf-7(e1372)* and *daf-9(m540)* mutants). Where possible, we assayed controls from treated wildtype mothers which had been previously mated with mutant males, such that the genotypes of the heterozygous progeny compared between control and treated conditions were identical. For *daf-7(e1372)* mutants, this was not possible due to the severe mating defect of males. For this strain, heterozygous progeny of mutant mothers were compared to homozygous wildtype controls, as noted in the main text. Additionally, for X-linked genes that were thus assayed (*daf-3, daf-12* and *odr-1*), we note that although the genotypes of the assayed hermaphrodites were identical across conditions, in this case they grew alongside male siblings of different genotypes (wildtype male siblings in the case of wildtype mothers but hemizygous mutant male siblings in the case of mutant mothers).

Wildtype males used were of the strain BCN2080 (Semple et al. 2012), which carries a singlecopy *pmyo-2::mCherry* transgene. The bright, stable pharyngeal fluorescence of the transgene was used as a paternity marker. Where mutant males were used for control crosses, the mutant strain was first crossed to BCN2080 to generate a new mutant strain with the paternity marker. Parental generation hermaphrodites were mated from young adulthood to *pmyo-2::mCherry* males at a ratio of 2-3 males per hermaphrodite, with 80-120 hermaphrodites per 60 mm plate per strain. After around 6h of mating, hermaphrodites were picked manually onto pheromone or control plates. Progeny could then be collected 18-20h later and transferred to a fresh plate for development and assay. Crosses thus performed normally yielded 100% cross progeny at the point of larvae collection. However the fluorescent paternity marker allowed for the removal of any contaminating self-fertilised progeny prior to assay. Male siblings of the assayed hermaphrodites were left on the plates and ignored during the assay - only hermaphrodites were scored.

*daf-7(e1372)* and *daf-9(m540)* hermaphrodites frequently enter dauer, leading to problematic heterogeneity in maternal population age. To bypass dauer and ensure a more developmentally synchronized maternal population, prior to crossing at the young adult stage we raised *daf-7* and *daf-9* mutant hermaphrodites from hatching on plates with *E. coli* OP50 supplemented with 100 nM (25S)-Δ7-dafachronic acid (Abcam, product ab144236) immediately prior to seeding. We confirmed that maternal dafachronic acid supplementation does not affect progeny soma-germline heterochrony (see Fig S6i).

### Heterozygous selfing parent experiments to assay mutation effect in the progeny generation

For *daf-3* and *daf-12* mutations we performed experiments where heterozygous parents were exposed to control- or pheromone-treated plates. In order to mark the wildtype chromosome, we identified strains from *wormbuilder.org* that contained a bright stable fluorescent marker integrated at a genetic location as close as possible to the relevant mutation. In this way recombination frequencies between the fluorescent marker and the mutant alleles would be negligible, allowing us to use the fluorescent marker as a proxy for the wildtype allele. For *daf-12* we used strain EG7992, marker genetic position X: 1.87, 0.5 cM from *daf-12* (estimated 0.50 % recombination rate). For *daf-3*, we used strain EG7972, marker genetic position X:-18.91, 0.53 cM from daf-3 (estimated 0.53% recombination rate). Background *unc-119* mutations were crossed out of these strains prior to use in these experiments.

Males carrying the wildtype allele and fluorescent marker were crossed to mutant hermaphrodites. Fluorescent heterozygous virgin hermaphrodite F1 progeny were picked to a fresh plate at L4 and moved to control- or pheromone-treated plate as young adults. Around 18-24h later progeny were collected by hatching and moved to a fresh plate. At the L4 stage, around 100 non-fluorescent homozygous mutant progeny were picked onto a fresh plate, while around 100 fluorescent heterozygous or homozygous wildtype siblings were picked onto a second fresh plate. Somatic and germline development were then assayed in these worms.

### *Experiment with* fog-2 *feminized worms to assay maternal or paternal transmission*

In order to investigate whether exposure to pheromone could influence progeny phenotypes via the paternal line, synchronized populations of N2 males were established by a 2 h egg-lay with male-fertilized hermaphrodites. Around 200 young adult males were transferred by picking to previously prepared pheromone or control plates and were incubated on these plates for 24 h. Following incubation with pheromone, males were washed off these plates and transferred by pipetting to a plate with around 100 young adult spermless *fog-2(q71)* mutant hermaphrodites that had been synchronized by egg-laying. Males had been removed from *fog-2* mutant plates prior to the L4/young adult molt to ensure that hermaphrodites remained virgins. N2 males were allowed to mate with *fog-2* hermaphrodites for 2 h before being removed. Around 18 h later hermaphrodites and hatched larvae were washed from the plates and larvae hatching during a 1-2h interval were collected and transferred to fresh plates for development and assay. As *fog-2* hermaphrodites are unable to self-fertilize, all progeny were sired by the pheromone- or control-treated N2 males. All progeny were heterozygous for the *fog-2* mutation and thus hermaphrodites were capable of self-fertilization. In the reciprocal experiment conducted in parallel to confirm transmission via the female line, *fog-2* mutant hermaphrodites were mated for 2 h by N2 males on a single plate. Following mating, fertilized hermaphrodites were manually transferred by picking to pheromone or control plates before progeny were collected for assay around 18-24h later.

### Timing of initiation of soma/germline post-embryonic development

Division of the two primordial germ cells (PGCs) was assayed using the germline-expressed *mCherry::H2B* fusion in strains BCN9111 and BCN9112, derived from strain EG6787. Larvae with two germ cell nuclei were counted as undivided, larvae with three or more clearly separated germ cell nuclei were scored as divided. As previously reported (Butuči et al. 2015), we observed an asymmetric initial division of the PGCs, with one cell dividing before the other. H cell divisions were assayed using a *SCMp:gfp* seam cell reporter in strain BCN9112, derived from strain JR667. H cells are the three anterior-most pairs of seam cells (H0, H1 and H2) and are the last group of seam cells to undergo initial divisions in L1 larvae. We chose to assay H cells instead of the earlier-dividing and more posterior V cells because they divide at a similar time to the PGCs in control worms. Worms were scored as divided when at least one of the six H cells had divided. The division of the single post-embryonic mesoblast, the M cell, was assayed with an *hlh-8:gfp* fusion in strains BCN9111, derived from strain PD4666.

To assay the impact of pheromones on timings of initial post-embryonic divisions in fed larvae, we first divided a synchronized cohort of day 1 adults between plates conditioned with crude pheromone or control medium. Around 24 hours later, we synchronized a population of worms by washing all adults and larvae off these plates with M9 buffer and allowing embryos to hatch. After 1 hour, the newly hatched larvae were collected in M9 buffer. Each treatment group was immediately divided between 4 fresh NGM plates seeded with OP50 to continue development. These worms were not starved at any point, as they hatch on a plate with food and are rapidly replated on food after collection. Hatching and development were conducted at 25 °C for these assays. From around 3.5 h after initial collection, larvae were periodically assayed using epifluorescence microscopy conducted with a DMI6000B inverted microscope (Leica) fitted with an ORCAFlash4.0 V2 Digital CMOS camera (Hamamatsu). To assay, larvae were collected with M9 buffer into a plastic 15 ml tube from one plate each per treatment group. The tube was topped up with M9 + 0.01% Triton-X before worms were spun down at 200 *g* for 1 min. Supernatant was aspirated to 100 μl. 1 M NaN_3_ was then added to a final concentration of 50 mM in order to rapidly anaesthetize the worms and halt development. Anaesthetized worms were transferred by pipetting into a 96 well plate. Worms were observed through the camera with live view using MetaMorph software (Leica). At each time point, the number of worms with relevant cells either divided or undivided were counted manually. Note as examination of germ cells particularly required inspection in the Z plane these assays were unsuitable for automated counting. Time points were taken every 1-1.5 h until more than 50% of the worms had undergone cell division. As described above for staging soma and germline development, count data was then modelled using a binomial generalized linear model, and the time at which half of the population had undergone the developmental transition (T_50_) was estimated using the function ‘dose.p’ of the library ‘MASS’ in R. 95% confidence intervals were estimated by multiplying the standard error obtained by the ‘dose.p’ function by 1.96. The impact of pheromone on a particular cell division was calculated by subtracting the relevant T_50_ from control group from the T_50_ of the pheromone treated group. 95% confidence intervals were then estimated as described above for Δ soma-germline values.

### Brightfield and fluorescence microscopy

Brightfield microscopy to measure the length of L1 larvae at hatching and fluorescence microscopy to measure autofluorescence and VIT-2::GFP fluorescence was conducted as described previously in (Perez et al. 2017).

### Single worm luminometry to assay development

Developmental timings of single worms were determined by luminometry as in (Olmedo et al. 2015). Briefly, parents were exposed to pheromone- or control-conditioned plates for ~24h before single embryos were transferred into a well of a white, flat-bottom, 96-well plate by manual picking with an eyelash. Each well contained 200 μl of S-basal media with 10 g l^−1^ *E. coli* OP50 (wet weight) and 100 μM D-Luciferin. Plates were sealed with a gas-permeable cover (Breathe Easier, Diversified Biotech). We measured luminescence (Berthold Centro LB960 XS3) for 1 sec, at 5-min intervals. We performed all experiments inside temperature-controlled incubators (Panasonic MIR-154). Temperature was monitored using a Thermochron iButton (Maxim Integrated).

The raw data from the luminometer were analyzed as described in (Olmedo et al. 2015). p-values on representative data shown in figures come from unpaired *t* tests on only the data shown. p-values cited in the text come from combined analysis of all replicates and batches by linear mixed model analysis. For combined analysis, data were first normalized by subtracting the corresponding mean statistic of the control group from the data for each individual worm to allow effects to be analyzed across replicates. A linear mixed model was fitted using the lme4 package in R, fitting the reported outcome by pheromone treatment and hatching time as explanatory variables and replicate as a random effect. To obtain p-values and test for significance an ANOVA was performed comparing this model with a null model lacking pheromone treatment as an explanatory variable. 9 tests in total were performed, testing L1 to adult, L1 molt to adult, L1, L2, L3, L4, all molts, L2 to adult and L2+L3+L4. Reported p-values were thus Bonferroni-adjusted for 9 tests. Times for L1 to adult, L1 and all molts were significantly affected by pheromone treatment.

---

**Table S1.** List of strains used in this study.

| Strain name | Genotype | Notes |
| --- | --- | --- |
| AA86 | *daf-12(rh61rh411)* X |  |
| BCN2080 | *pBCN26[Prpl28:PuroR:Prpl16:NeoR operon, Pmyo2::mCherry]*, integrated as single copy | Paternity marker strain |
| BCN9071 | *vit-2::gfp* X | CRISPR knock-in |
| BCN9105 | *daf-12(rh61rh411)* X;<br>*pBCN26[Prpl28:PuroR:Prpl16:NeoR operon, Pmyo2::mCherry]* | Paternity marker strain |
| BCN9111 | *oxSi487 [mex-5p::mCherry::H2B::tbb-2 3'UTR::gpd-2 operon::GFP::H2B::cye-1 3'UTR + unc-119(+)]* II. *ayIs6* X [pBH47.70 (*hlh-8::GFP* fusion)] | Derived from crossing strains PD4666 and EG6787 |
| BCN9112 | *oxSi487 [mex-5p::mCherry::H2B::tbb-2 3'UTR::gpd-2 operon::GFP::H2B::cye-1 3'UTR + unc-119(+)]* II. *wIs51 V [SCMp::GFP + unc-119(+)]* | Derived from crossing strains JR667 and EG6787 |
| BCN9119 | *tax-4(ok3771)* III;<br>*pBCN26[Prpl28:PuroR:Prpl16:NeoR operon, Pmyo2::mCherry]* | Paternity marker strain, derived from crossing strains BCN2080 and VC3113 |
| BCN9120 | *tax-2(ok3403)* I;<br>*pBCN26[Prpl28:PuroR:Prpl16:NeoR operon, Pmyo2::mCherry]* | Paternity marker strain, derived from crossing strains BCN2080 and RB2464 |
| BCN9121 | *tax-2(p694)* I;<br>*pBCN26[Prpl28:PuroR:Prpl16:NeoR operon, Pmyo2::mCherry]* | Paternity marker strain, derived from crossing strains BCN2080 and PR694 |
| BCN9122 | *odr-1(n1936)* X;<br>*pBCN26[Prpl28:PuroR:Prpl16:NeoR operon, Pmyo2::mCherry]* | Paternity marker strain, derived from crossing strains BCN2080 and CX2065 |
| BCN9123 | *daf-11(m47)* V;<br>*pBCN26[Prpl28:PuroR:Prpl16:NeoR operon, Pmyo2::mCherry]* | Paternity marker strain, derived from crossing strains BCN2080 and DR47 |
| BCN9125 | *daf-3(e1376) X*;<br>*pBCN26[Prpl28:PuroR:Prpl16:NeoR*<br>*operon, Pmyo2::mCherry]* | Paternity marker<br>strain |
| CB1372 | *daf-7(e1372) III* |  |
| CB1376 | *daf-3(e1376) X* |  |
| CB4108 | *fog-2(q71) V* |  |
| CX2065 | *odr-1(n1936) X* |  |
| DR2281 | *daf-9(m540) X* |  |
| DR47 | *daf-11(m47) V* |  |
| DR476 | *daf-22(m130) II* |  |
| EG7972 | *unc-119(ed3) III*; *oxTi673 X* | Chromosomal marker<br>for wildtype *daf-3* |
| EG7992 | *unc-119(ed3) III*; *oxTi545 X* | Chromosomal marker<br>for wildtype *daf-12* |
| JN1715 | *peIs1715 [str-1p::mCasp-1 + unc-122p::GFP]* | AWB ablation |
| MOL006 | *sevls1[Psur-5::luc+::gfp] X* | For single-worm<br>luminometry |
| N2 | - | wildtype strain for<br>this study |
| PR694 | *tax-2(p694) I* |  |
| PS6025 | *qrIs2 [sra-9::mCasp1]* | ASK neuron ablation |
| PY7502 | *oyIs85 [ceh-36p::TU#813 + ceh-36p::TU#814 + srtx-1p::GFP + unc-122p::DsRed]* | AWC neuron ablation |
| PY7505 | *oyIs84 [gpa-4p::TU#813 + gcy-27p::TU#814 + gcy-27p::GFP + unc-122p::DsRed]* | ASI ablation |
| RB2464 | *tax-2(ok3403) I* |  |
| VC3113 | *tax-4(ok3771) III* |  |
| VS8 | *dhs-28 (hj8) X* |  |
| ZD763 | *mgIs40[daf-28p::nls-GFP]*;<br>*jxEx100[trx-1::ICE + ofm-1::gfp]* | ASJ ablation. Gift of<br>Dennis Kim. |

**Figure S1.**
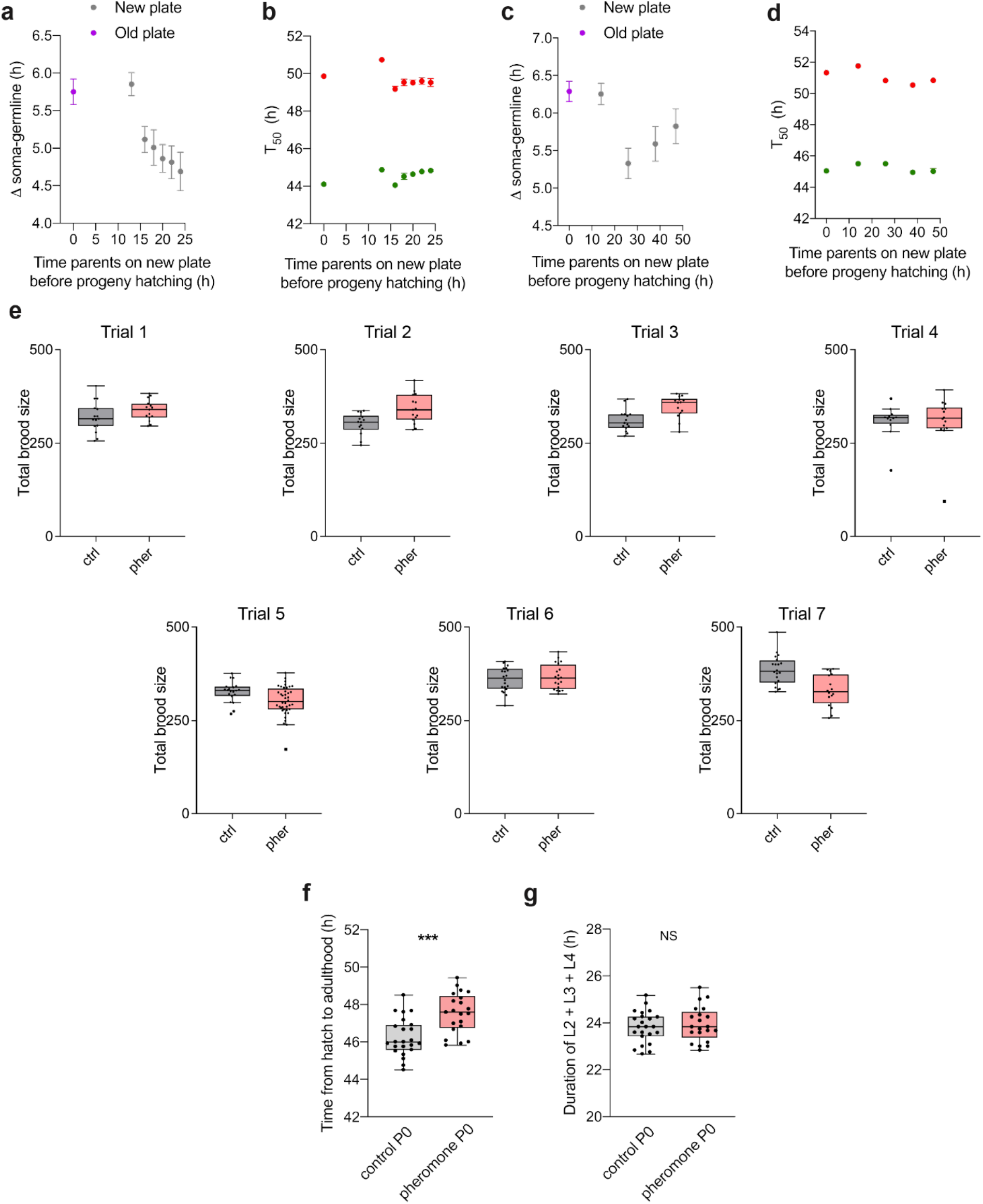
Progeny germline acceleration peaks around 1 day after parents are moved to fresh plate, while pheromone exposure does not affect brood size but delays somatic development by extending the L1 stage. **a)** Time between somatic and germline maturity for worms whose parents were left on the conditioned plate (grey) or shifted to a fresh plate at various intervals prior to progeny hatching (pink). T_50_ measurements shown in **b**. n = 141-313. **b)** Timings of the L4/YA molt (green) and first embryo (red) for data shown in **a**. **c)** Time between somatic and germline maturity for worms whose parents were left on the old plate (grey) or shifted to a new plate at various intervals prior to progeny hatching (pink). T_50_ measurements shown in **c**. n = 137-253. **d)** Timings of the L4/YA molt (green) and first embryo (red) for data shown in **c**. **e)** Total brood size data for worms whose parents were exposed to plates conditioned with control extract or crude pheromone extract across 7 trials. n = 15, 15 (Trial 1), 15, 15 (Trial 2), 15, 15 (Trial 3), 14, 15 (Trial 4), 20, 40 (Trial 5), 20, 20 (Trial 6), 20, 16 (Trial 7). **f)** Boxplot showing time from hatch to adulthood for single worms in liquid culture whose parents were exposed to control (grey) or pheromone-conditioned (pink) plates. n= 23, 21. **g)** Boxplot showing the combined duration of the L2, L3 and L4 larval stages for single worms whose parents were exposed to control (grey) or pheromone-conditioned (pink) plates. n= 23, 21. Panels **f** and **g** show data from a single representative replicate. Significance levels shown in panels **f** and **g** are from unpaired t tests on the data shown, whereas p values cited in main text are from linear mixed model analysis of 3 replicates pooled. Error bars in panels **a**-**d** show 95% confidence interval (may not be visible when error is small). Boxplots show Tukey whiskers with data points overlaid. Ctrl, control-conditioned plates. Pher, crude pheromone conditioned plates. P0, parental generation. NS, not significant. ***, p < 0.001.

**Figure S2.**
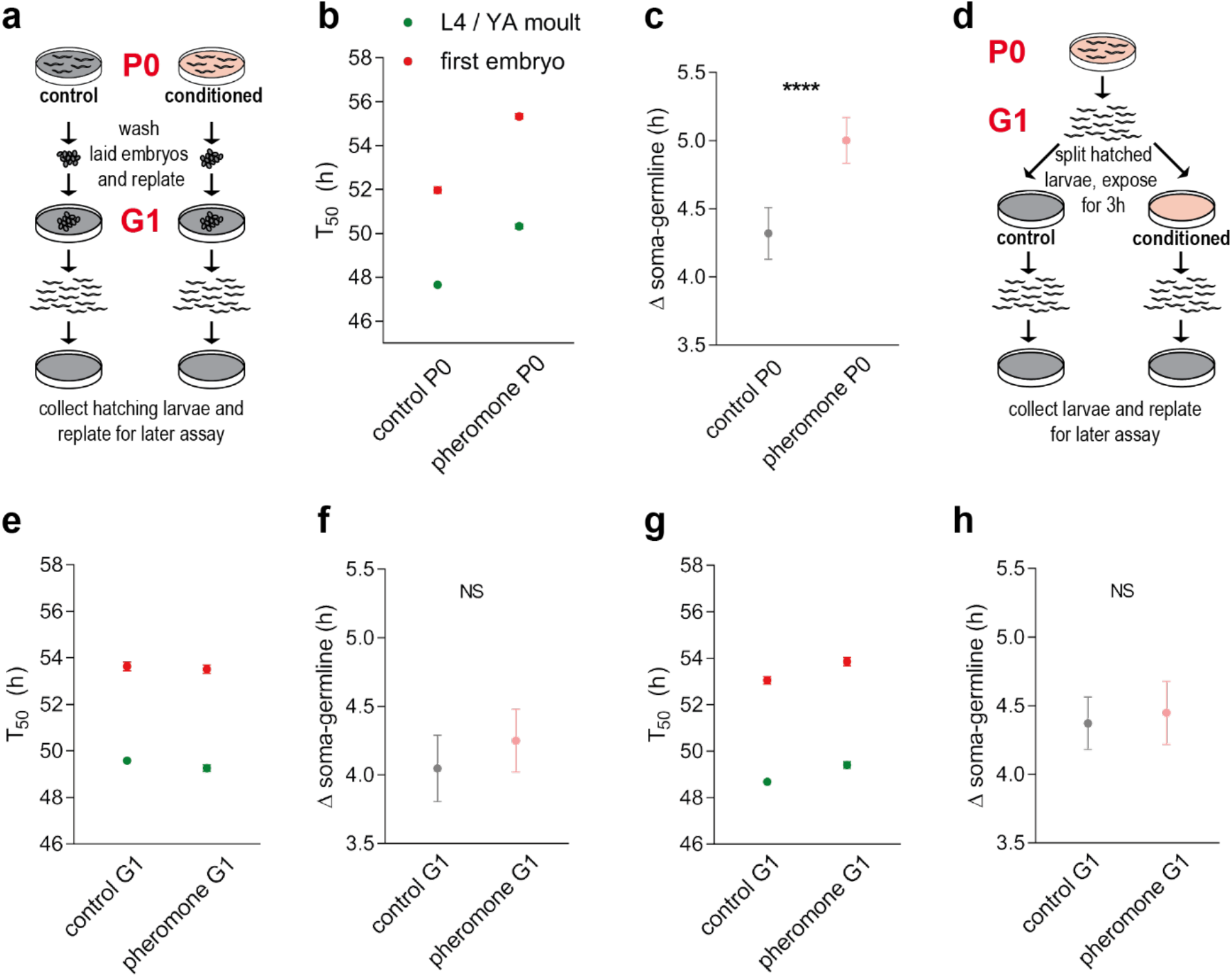
Germline delay in progeny is due to parental worm pheromone exposure and not to direct exposure within the same generation. **a)** Experimental schematic demonstrating the harvest of embryos laid by parents on control or pheromone-conditioned plates. Embryos are briefly bleached and thoroughly washed before replating on a fresh unconditioned plate. Hatchers on fresh unconditioned plates are collected a few hours later to assay developmental timings. **b)** Timings of the L4/YA molt (green) and first embryo (red) for worms hatched on fresh plates from thoroughly washed embryos laid by control or pheromone-exposed parents. n = 156-358. **c)** The time between somatic maturity and germline maturity for worms whose parents were moved to plates conditioned with control extract or crude pheromone extract, derived from the T_50_ measurements shown in panel **b**. **d)** Experimental schematic demonstrating the early 3h transient exposure of hatchers from a single plate to control extract or crude pheromone extract. **e)** Timings of the L4/YA molt (green) and first embryo (red) for worms exposed transiently to control or pheromone-conditioned plates. n = 178-306. **f)** The time between somatic maturity and germline maturity for worms exposed transiently to control or pheromone-conditioned plates, derived from the T_50_ measurements shown in panel **e**. **g)** Timings of the L4/YA molt (green) and first embryo (red) for worms exposed continuously throughout development to control or pheromone-conditioned plates. n = 176-275. **h)** The time between somatic maturity and germline maturity for worms exposed continuously throughout development to control or pheromone-conditioned plates, derived from the T_50_ measurements shown in panel **g**. All experiments were replicated at least 3 times independently. Error bars show 95% confidence interval (may not be visible when error is small). All panels show representative data from a single trial. P0, parental generation. G1, first progeny generation. YA, young adult. NS, not significant. **\*\*\*\***, p < 0.0001.

**Figure S3.**
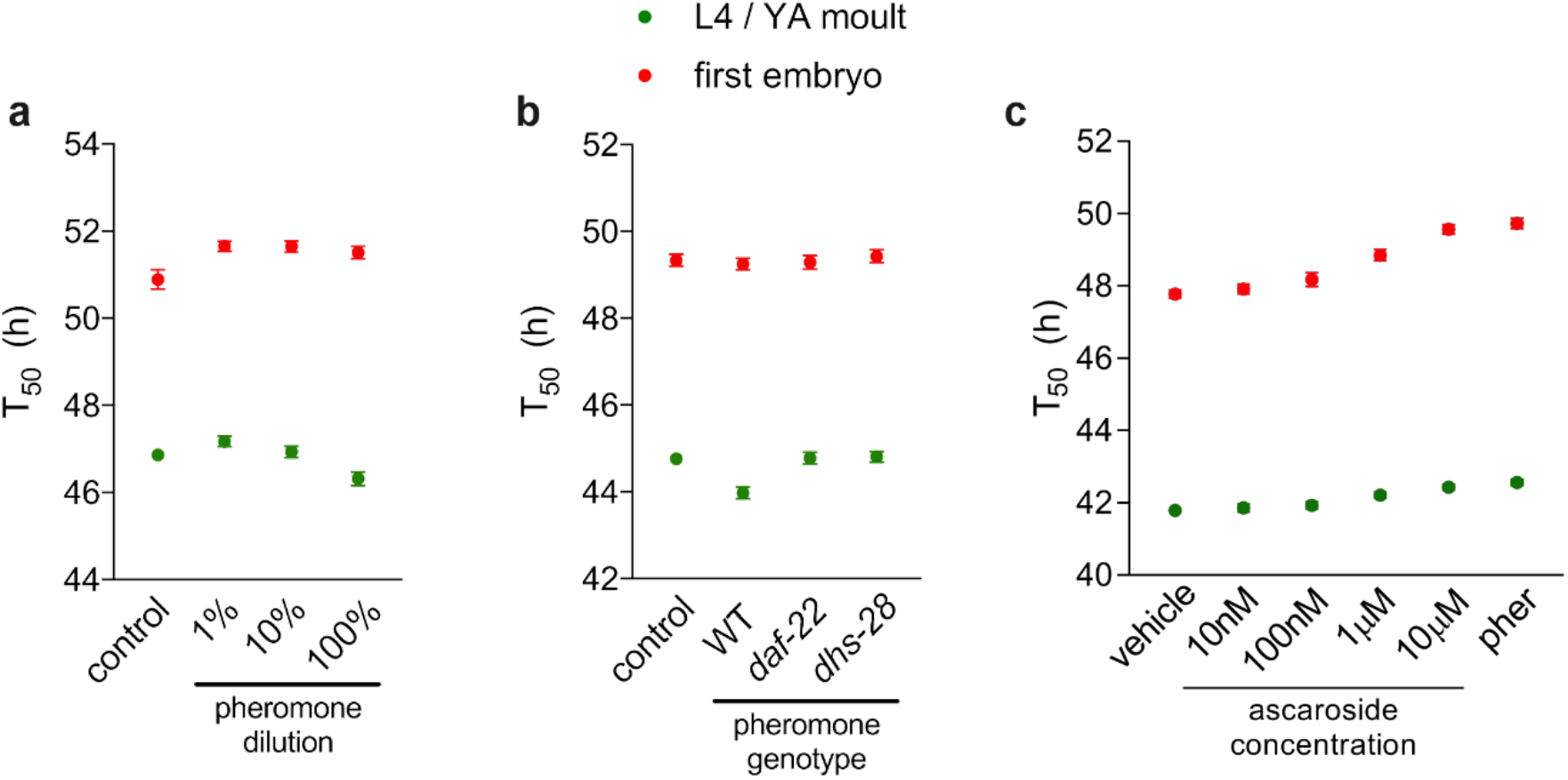
The effect of crude pheromone extract is dose-dependent and is likely to be mediated in part by major ascarosides (related to Figure 2). **a)** Timings of the L4/YA molt (green) and first embryo (red) for worms whose parents were exposed to various dilutions of crude pheromone extract. n = 214-429. **b)** Timings of the L4/YA molt (green) and first embryo (red) for worms whose parents were exposed to control plates, wildtype pheromone extract, or *daf-22* or *dhs-28* mutant pheromone extract. n = 137-283. **c)** Timings of the L4/YA molt (green) and first embryo (red) for worms whose parents were exposed to vehicle-only control plates or various concentrations of an equimolar blend of synthetic major ascarosides #2 and #3. n = 191-337. All experiments were replicated at least 2 times independently. Error bars show 95% confidence interval (may not be visible when error is small). All panels show representative data from a single trial.

**Figure S4.**
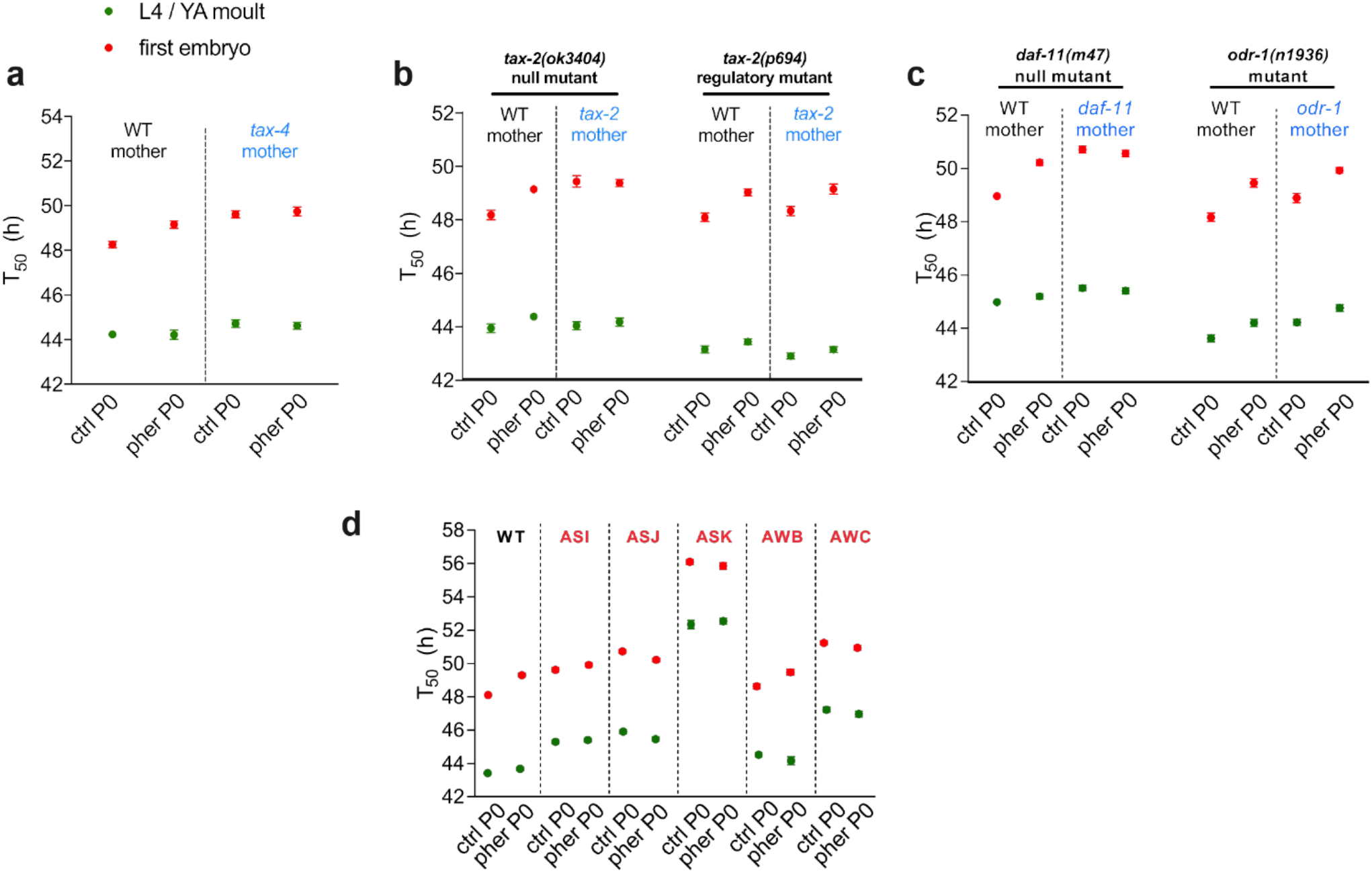
The TAX-2/TAX-4 cyclic nucleotide-gated channels and the upstream guanylyl cyclase DAF-11 are required in parental neurons for progeny response to exposure of parents to pheromone (related to Figure 3). **a)** Timings of the L4/YA molt (green) and first embryo (red) for *tax-4* heterozygous worms whose mothers were wildtype hermaphrodites (left) or homozygous *tax-4(ok3771)* mutant hermaphrodites (right) exposed to control plates or wildtype pheromone extract. n = 81 - 210. **b)** Timings of the L4/YA molt (green) and first embryo (red) for worms heterozygous for either the *tax-2(ok3404)* null allele (left half) or *tax-2(p694)* regulatory mutant (right half) whose mothers were wildtype hermaphrodites (left of dashed line) or homozygous *tax-4(ok3771)* mutant hermaphrodites (right of dashed line) exposed to control plates or wildtype pheromone extract. n = 129-250. **c)** Timings of the L4/YA molt (green) and first embryo (red) for worms heterozygous for mutant alleles of the receptor-bound guanylyl cyclases *daf-11(m47)* (left half) or *odr-1(n136)* (right half) whose mothers were wildtype hermaphrodites (left of dashed line) or homozygous mutant hermaphrodites (right of dashed line) exposed to control plates or wildtype pheromone extract. n = 57-271. **d)** Timings of the L4/YA molt (green) and first embryo (red) for worms with genetic ablations of the ASI, ASJ, ASK, AWB or AWC chemosensory neurons exposed to control plates or wildtype pheromone extract. n = 125-264. All experiments were replicated at least 2 times independently. All panels show representative data from a single trial. All data shown in a single panel are from the same trial. P0, parental generation. WT, wildtype. Ctrl, control-conditioned plates. Pher, crude pheromone conditioned plates. Error bars in panels show 95% confidence interval (may not be visible when error is small).

**Figure S5.**
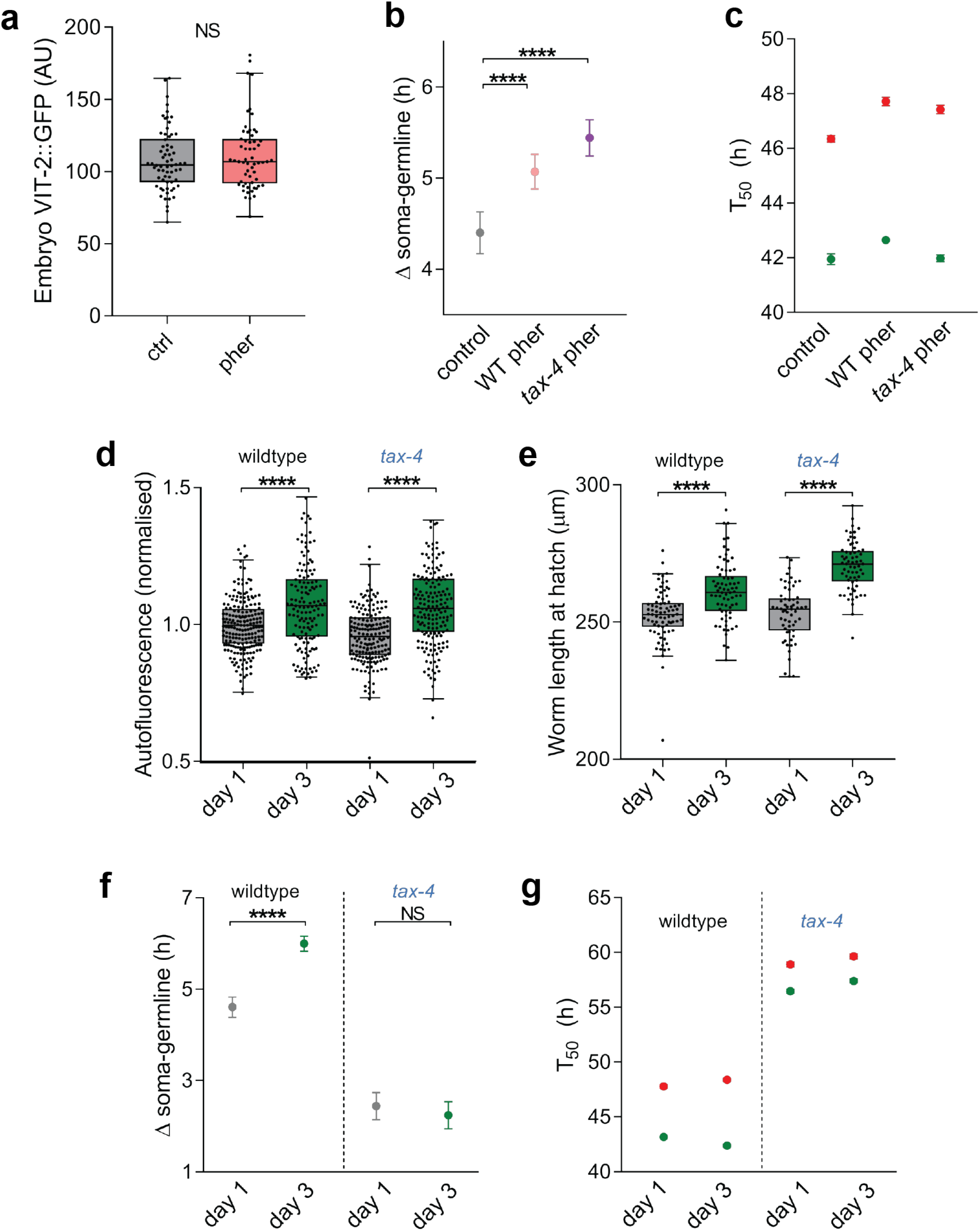
Pheromone-insensitive *tax-4* mutants do not show increasing relative germline delay with maternal age. **a)** Boxplot showing total fluorescence of a yolk protein/vitellogenin fusion reporter, VIT-2::GFP, in early embryos from parents treated with control (grey) or pheromone-conditioned (pink) plates. **b)** The time between somatic and germline maturity for worms whose parents were exposed to control (grey), wildtype pheromone-conditioned (pink) or *tax-4* pheromone-conditioned (purple) plates. T_50_ measurements shown in **c**. **c)** Timings of the L4/YA molt (green) and first embryo (red) for worms whose parents were exposed to control, wildtype pheromone-conditioned or *tax-4* pheromone-conditioned plates. **d)** Boxplot showing autofluorescence, a proxy for vitellogenin content, in early embryos from day 1 or 3 wildtype mothers (left of dashed line) or *tax-4* mothers (right of dashed line). **e)** Boxplot showing length of L1 larvae at hatch from day 1, 2 or 3 wildtype mothers (left of dashed line) or *tax-4* mothers (right of dashed line). **f)** The time between somatic and germline maturity for worms whose parents were wildtype(left of dashed line) or *tax-4* mothers (right of dashed line) on day 1 (grey) or day 3 (green) of adulthood. **g)** Timings of the L4/YA molt (green) and first embryo (red) for worms whose parents were wildtype(left of dashed line) or *tax-4* mothers (right of dashed line) on day 1 or day 3 of adulthood. All experiments were repeated at least 3 times independently. All data show a single representative trial. Data from panel **d** comes from 3 replicates pooled and analysed together. Data are normalized to the mean of wildtype day 1 in each trial to allow plotting together. Data from panels **e**, **f** and **g** come from the same trial. Boxplots show Tukey whiskers with data points overlaid. Ctrl, control-conditioned plates. Pher, crude pheromone conditioned plates. WT, wildtype. NS, not significant. ****, p < 0.0001.

**Figure S6.**
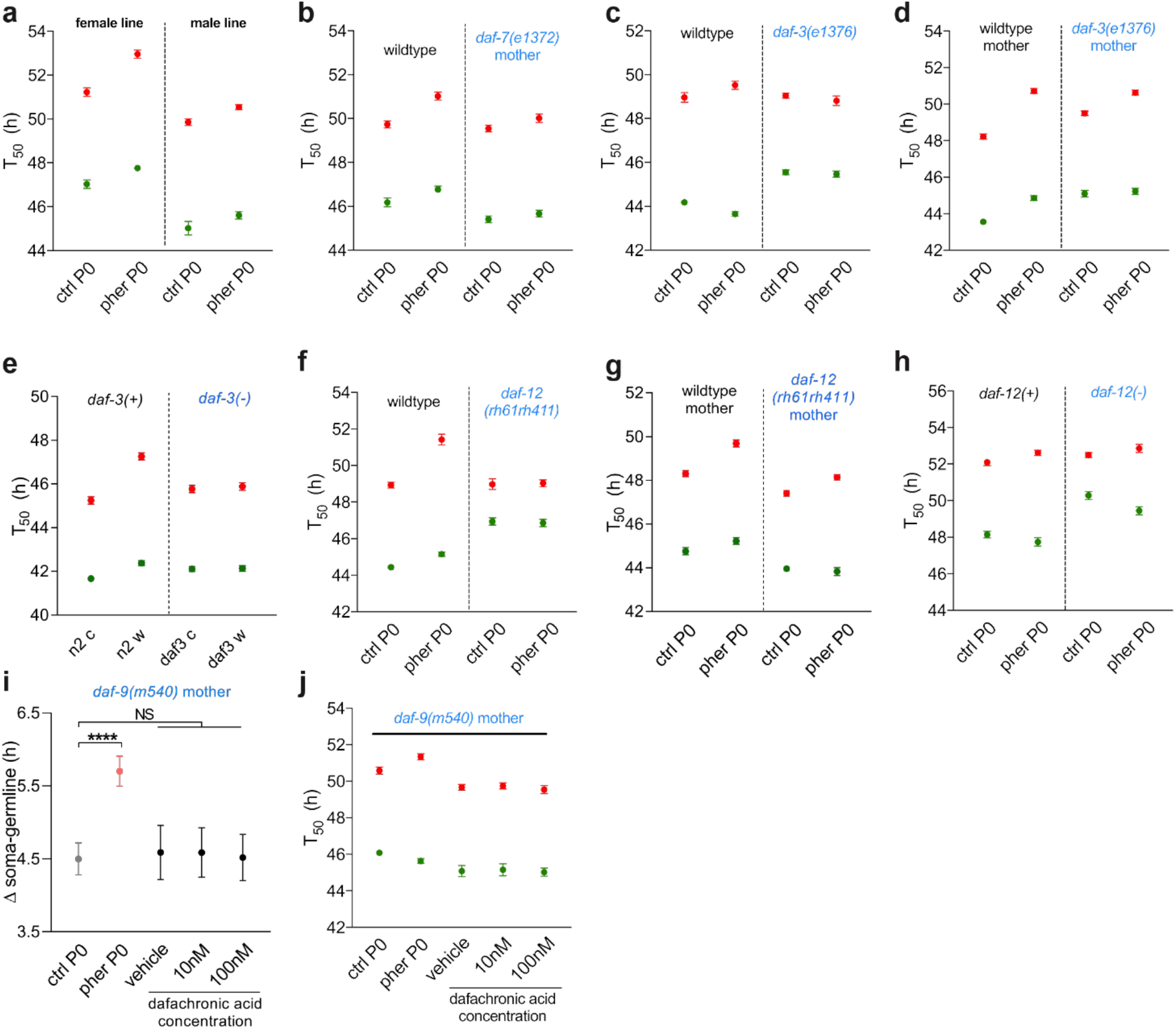
Effects of parental pheromone exposure are inherited via the oocyte and require DAF-7/TGF-β in parental neurons and the Co-SMAD DAF-3 and nuclear hormone receptor DAF-12 in progeny (related to Figure 5). **a)** Timings of the L4/YA molt (green) and first embryo (red) for male-sired worms whose mothers (left of dashed line) or fathers (right of dashed line) were exposed to control plates or crude pheromone. n = 94-154. **b)** Timings of the L4/YA molt (green) and first embryo (red) for male-sired wildtype worms (left of dashed line) or male-sired heterozygous progeny of *daf-7(e1372)* mutant mothers (right of dashed line) whose mothers were exposed to control plates or crude pheromone. n = 102-200. **c)** Timings of the L4/YA molt (green) and first embryo (red) for wildtype worms (left of dashed line) or *daf-3(e1376)* mutant worms (right of dashed line) whose parents were exposed to control plates or crude pheromone. n = 120-242. **d)** Timings of the L4/YA molt (green) and first embryo (red) for *daf-3* heterozygous progeny whose wildtype mothers (left of dashed line) or *daf-3(e1376)* mutant mothers (right of dashed line) were exposed to control plates or crude pheromone. n = 73-233. **e)** Timings of the L4/YA molt (green) and first embryo (red) for *daf-3(+)* (left of dashed line) or homozygous *daf-3(e1376)* mutant (right of dashed line) progeny whose *daf-3* heterozygous mothers were exposed to control plates or crude pheromone. n = 85-135. **f)** Timings of the L4/YA molt (green) and first embryo (red) for wildtype worms (left of dashed line) or *daf-12(rh61rh411)* mutant worms (right of dashed line) whose parents were exposed to control plates or crude pheromone. n = 102-208. **g)** Timings of the L4/YA molt (green) and first embryo (red) for *daf-12* heterozygous progeny whose wildtype mothers (left of dashed line) or *daf-12(rh61rh411)* mutant mothers (right of dashed line) were exposed to control plates (grey) or crude pheromone (pink). n = 63-181. **h)** Timings of the L4/YA molt (green) and first embryo (red) for *daf-12(+)* (left of dashed line) or homozygous *daf-12(rh61rh411)* mutant (right of dashed line) progeny whose *daf-12* heterozygous mothers were exposed to control plates or crude pheromone. n = 67-194. **i)** Time between somatic and germline maturity for male-sired *daf-9* heterozygous progeny of *daf-9(m540)* mutant mothers exposed to control plates (grey), crude pheromone (pink) or various concentrations of (25S)-Δ7-dafachronic acid (black). T_50_ measurements shown in **j**. n = n = 67-218. **j)** Timings of the L4/YA molt (green) and first embryo (red) for for male-sired *daf-9* heterozygous progeny of *daf-9(m540)* mutant mothers exposed to control plates, crude pheromone or various concentrations of (25S)-Δ7-dafachronic acid. n = 67-218. All experiments were replicated at least 2 times independently. All panels show representative data from a single trial. All data within a panel comes from the same trial. Error bars in panels show 95% confidence interval. Pairwise tests are subject to Bonferroni corrections for multiple testing within each trial. Ctrl, control-conditioned plates. Pher, crude pheromone conditioned plates. P0, parental generation. NS, not significant. ****, p < 0.0001.

**Figure S7.**
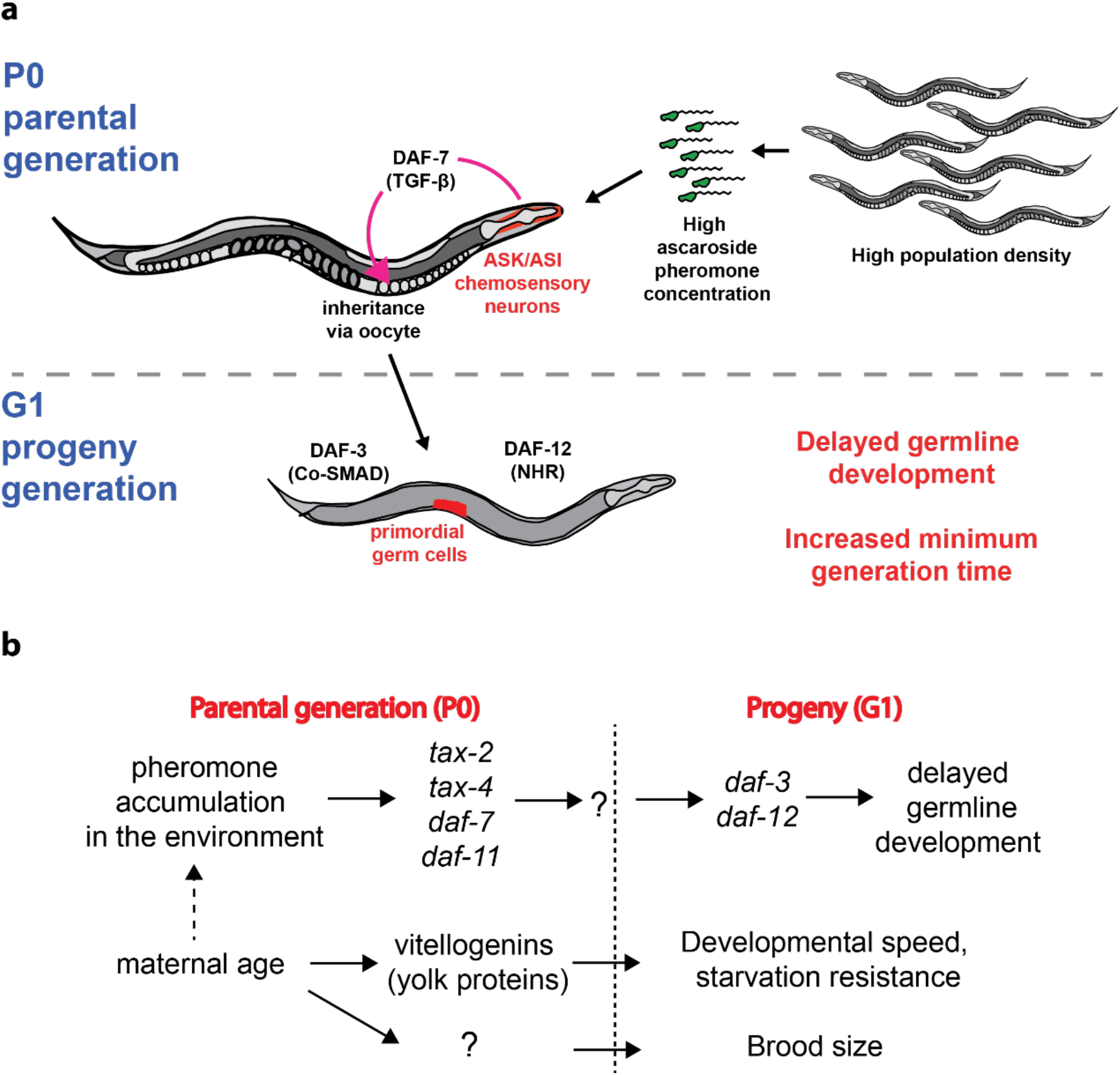
Summary of the reported intergenerational effects on phenotypic variation in *C. elegans*. **a)** Our results are consistent with a model whereby population density, communicated by ascaroside pheromone concentration, is perceived in the parent’s chemosensory neurons and transmitted via the TGF-β ligand DAF-7. The signal is inherited via the oocyte, crossing the Weismann barrier and delaying the initial divisions of the primordial germ cells of progeny. This leads to a delay in germline maturity in adulthood and an increased generation time. Germline delay requires DAF-3/Co-SMAD and DAF-12/NHR. **b)** Maternal age influences progeny physiology via at least three inherited signals. Maternal age causes progeny germline delay directly via increased exposure to pheromone of older mothers. Maternal age determines size at hatching, developmental speed and resistance to L1 starvation via an increasing supply of the yolk protein vitellogenin to embryos. Maternal age also influences progeny brood size via an unknown mechanism.

## ACKNOWLEDGMENTS

We would like to thank Frank Schroeder for his kind gift of synthetic ascarosides and Dennis Kim for providing the ASJ neuron ablation strain. Thanks also to Thomas Wilhelm for help in the lab.

## AUTHOR CONTRIBUTIONS

MFP conceived and performed experiments, analyzed data and drafted the manuscript. MS and MF performed the experiments shown in Fig 2d and Fig S3c. SDV and MF performed the experiments shown in Fig 2a, Fig S1a-d and Fig S3a. AMC and MO performed the experiments shown in Fig 1g,h and Fig S1f,g and analyzed the data. MF discovered the effect, conceived and performed experiments, analyzed data and edited the manuscript. BL conceived experiments and edited the manuscript.

